# Functional genomics in a microbe that degrades and metabolizes PET plastic

**DOI:** 10.1101/2025.11.11.687616

**Authors:** Alexander Crits-Christoph, Julia Leung, Felipe-Andrés Piedra, Stephanie L. Brumwell, Victoria A. Sajtovich, Melanie B. Abrams, Ariela Esmurria, Shinyoung Clair Kang, Kerrin Mendler, Charlie Gilbert, Henry H. Lee, Nili Ostrov

## Abstract

*Piscinibacter sakaiensis* (formerly *Ideonella sakaiensis*) was the first bacterial species known to both completely degrade and assimilate polyethylene terephthalate (PET). However, the absence of efficient genetic tools has limited direct engineering of this organism, forcing most efforts to rely on heterologous expression of PET-degrading enzymes in model hosts. Here, we establish foundational genetic tools to engineer *P. sakaiensis*. We identify a functional plasmid origin of replication, multiple new selectable markers, and a transposon system for the strain. We use these tools to construct a genome-wide, barcoded transposon mutant library for pooled high-throughput functional screens. We apply this mutant library to growth on PET and identify metabolic and physiological genes that impact PET biodegradation. We also use this library to reveal mutants with improved DNA uptake for genome engineering. Together, these advances provide a platform for functional genomics in *P. sakaiensis* and positions this naturally evolved plastic-degrading bacterium as an engineerable chassis for synthetic biology and sustainable materials research.

## Introduction

Large-scale plastic production began in the middle of the 20th century. Since then, the industry and its waste have grown explosively^1,2^, while less than a tenth of plastics have been recycled once and less than a hundredth have been recycled more than once^1^. Each year, approximately 10 million metric tons of plastic waste reach aqueous environments, and the increase in waste has outpaced efforts to improve waste management^3^. The volume of waste, both current and projected, have prompted international concern, with the United Nations noting threats to ecosystems, fisheries, and human societies^4^.

Plastics persist in the environment and are recalcitrant to biological degradation, which is a benefit for the consumer, but a significant challenge for waste management^5^. Physical recycling of plastic currently cannot achieve circularity, as the recycled product is of lower quality than the virgin material, and current chemical recycling remains energy intensive and expensive^6,7^. With more carbon in the form of plastic entering the environment, interest in microbial-based plastic degradation has increased^8^. Efforts for enzymatic degradation of plastics such as polyethylene terephthalate (PET)^9^ also exist, and these enzymatic methods can require less stringent cleaning of the plastic substrate, reducing the cost of the recycling workflow as compared to typical mechanical processes^10,11^. However, harnessing a microbe capable of plastic degradation and conversion creates an opportunity to valorize this plastic waste directly, avoiding costly enzyme production and purification.

*Piscinibacter sakaiensis* 201-F6 (formerly *Ideonella sakaiensis*) was the first identified bacterial strain able to both efficiently and completely degrade low-crystallinity PET plastic^12^, the most abundant polyester plastic^1^. *P. sakaiensis* can utilize PET as a major carbon source, and recent research has shown media supplementation can further enhance the rate of PET degradation in the strain^13^. Nearly a decade has elapsed since its isolation, during which time substantial research efforts were dedicated to the *P. sakaiensis* PETase hydrolase that hydrolyzes PET polymers^14–19^. There has also been considerable research focused on the production of PETase enzymes in traditional laboratory microbial hosts such as *Escherichia coli* and *Pseudomonas*^20–26^, and heterologous expression of PETases in some environmental microbes^27^. However, there has been comparatively little investigation on engineering of native *P. sakaiensis*, despite this bacterium’s native ability to degrade amorphous PET efficiently. Homologous recombination has been used to edit the *P. sakaiensis* genome and identify the essentiality of two genes, the PETase and mono(2-hydroxyethyl) terephthalate hydrolase (MHETase), for growth on PET^28^. Yet, there are no reports to date for synthetic biology tools enabling high-throughput approaches for genetic engineering in *P. sakaiensis*.

## Results and Discussion

### Strain characterization

We performed a media and antibiotic screen to identify culture conditions and antibiotic susceptibility for *P. sakaiensis* 201-F6 (DSMZ 112585) (**Figure 1A-B**, **Supplementary Table 1**, **Supplementary Table 2**). We observed that *P. sakaiensis* reached the highest maximum optical density (OD_600_) in Horikoshi-1 medium (HMI) and Yeast Extract Malt Extract Dextrose (YEMED) (**Figure 1A**, **Supplementary Table 1**). However, the strain grew slightly faster in NBRC 802 medium, so we elected to use this media for further culturing. We found that the strain was susceptible to chloramphenicol (CAM), erythromycin (ERM), gentamicin (GEN), kanamycin (KAN), spectinomycin (SPEC), and tetracycline (TET), but not to carbenicillin (CARB) (**Figure 1B**, **Supplementary Table 2**).

**Figure 1.**
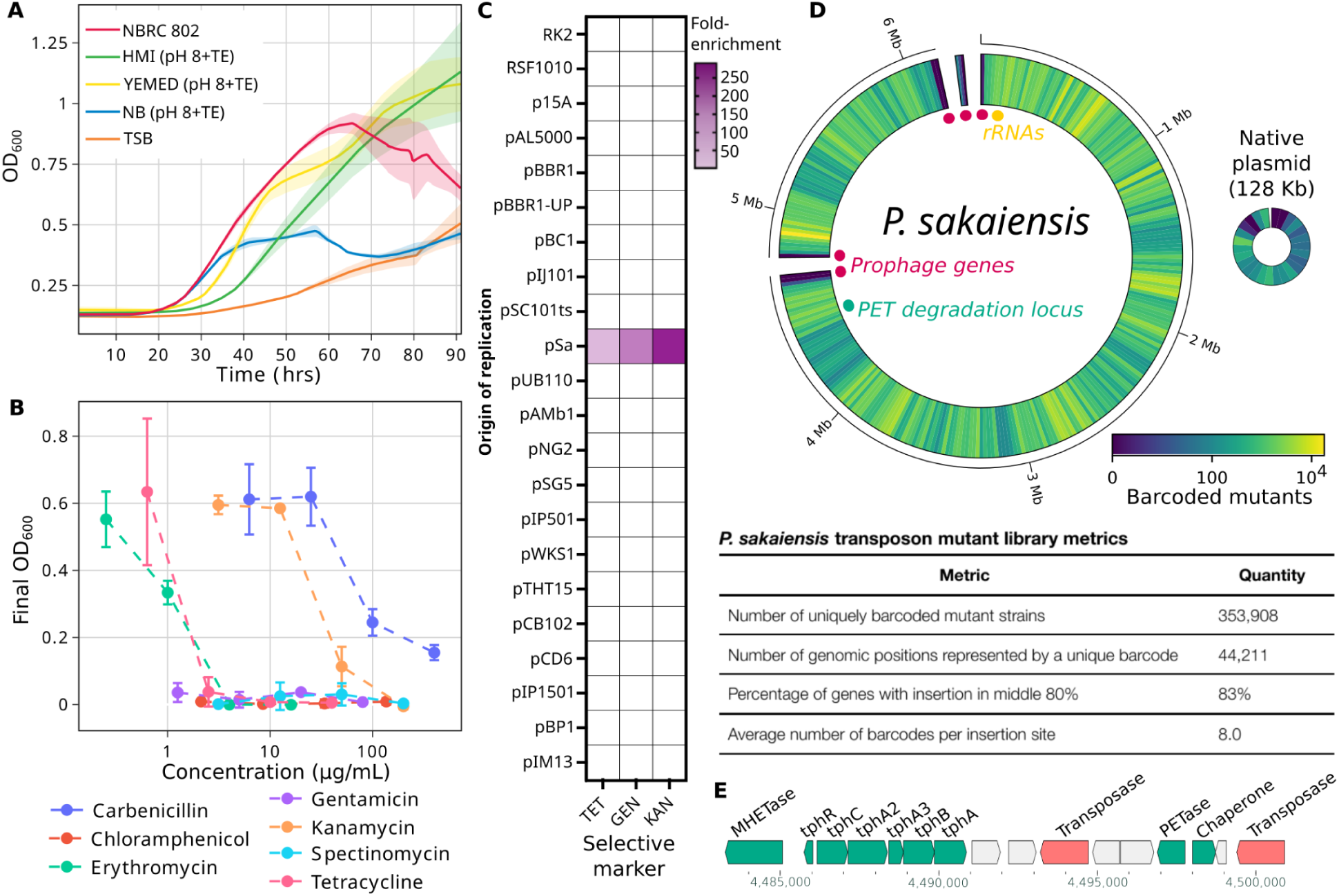
Strain growth and library characterization. **(A)** Growth curves of *P. sakaiensis* in five top-performing media. Full media details are found in **Supplementary Table 1**. **(B)** Antibiotic susceptibility of *P. sakaiensis* in liquid media. Error bars represent standard deviation. **(C)** POSSUM ORI-marker screen in *P. sakaiensis*. Heatmap of sequencing fold-enrichment of origins of replication in *P. sakaiensis* following conjugation with liquid selection with three POSSUM libraries with different selective markers. **(D)** Overview of the randomly-barcoded transposon mutant library in *P. sakaiensis*. Heatmap visualizing the number of unique barcoded mutants tiled across the three chromosomal scaffolds and plasmid scaffold of the *P. sakaiensis* genome. (**E**) Genomic visualization of the fully-assembled PET degradation locus, with contig coordinates ranging from the MHETase gene to the downstream PETase gene flanked by transposases. Genes predicted to be involved in PET degradation are highlighted in teal.

We assembled a genome of *P. sakaiensis* 201-F6 from Oxford Nanopore sequencing data that was 6.4 Mb in size and made up of four scaffolds: one scaffold represented a circular endogenous replicative 128 kb plasmid, and the other three were linked in the assembly graph, suggesting they together comprise the chromosome (NCBI BioSample accession: SAMN52661550). One 42 kb scaffold, found at twice the sequencing depth of the rest of the genome, represents a transposable prophage homologous to Bacteriophage Mu. This prophage accounts for the remaining two assembly breaks, as it is integrated at two positions separated by 1.3 Mb in the genome of the strain at time of sequencing, and may be recently active. In this genome the PETase gene is flanked by ISCR1 family transposases and is ∼6 kb downstream of the terephthalic acid (TPA) degradation operon and MHETase (**Figure 1E**), whereas in the originally reported genome the PETase was on a fragmented contig^12^. Genomic colocalization of the PETase with the MHETase provides evidence of a shared selective pressure^29^, presumably for PET degradation, and the presence of flanking transposases indicates that this selection may have occurred via a relatively recent transposition of the PETase gene.

To identify the potential impact of native restriction-modification (R-M) systems on DNA delivery, we characterized active methylation patterns with Nanopore detection of modified bases using MicrobeMod^30^ (**Supplementary Table 3**). We identified four active methylated sequence motifs and four R-M operons in the genome, suggesting that all four R-M systems are active (**Supplementary Table 3**). The observed methylated motif C**C**_5m_W**G**G matches the methylation activity of the Dcm methylase in *E. coli*, indicating compatibility with *dcm+* DNA in this strain. Recoding and removing occurrences of the other three methylated sequence motifs in genetic constructs could be particularly useful for improving transformation efficiency in this strain.

### Identification of a replicative plasmid for gene expression

We used the POSSUM toolkit^30,31^ to identify a functional origin of replication (ORI) and selectable markers (**Figure 1C**). Plasmid libraries with different antibiotic resistance markers, each harboring up to 23 ORIs, were delivered to *P. sakaiensis* by conjugation in solid media petri dish (GEN, KAN, TET) or 96-well plate (CAM, ERM, GEN, KAN, SPEC, TET) format. We obtained transconjugants for three libraries (GEN, KAN, and TET) using both formats, and sequencing identified pSa as a functional ORI (**Figure 1C**; **Supplementary Figure 1**). The pSa ORI was identified in all libraries and was validated by conjugation of an individual plasmid and whole genome sequencing. The pBBR1 ORI was identified in a single solid media sample, and was not validated (**Supplementary Figure 1**). This is the first report of a functional replicating plasmid for *P. sakaiensis*, with additional verification of both GEN and KAN as functional selectable markers in this species.

### Generation of a genome-wide barcoded transposon library

We next explored the use of transposons to develop a genome-wide mutant library. We chose the Himar-1 mariner transposon system which inserts at short 2 bp ‘TA’ motif sites^32^ and has been reported to work in multiple Gram-negative bacteria^33,34^. Although the *P. sakaiensis* genome is GC-rich (73%), there are still ∼193,000 TA sites in the genome, with 95% of all genes in the genome containing at least one TA site within the central 80% of the gene, suggesting that this transposon could be used for saturation transposon mutagenesis. We performed an initial pilot experiment by delivering the pMarC9-R6k transposon construct^35^ to *P. sakaiensis* via conjugation from *E. coli* BW29427. We obtained ∼100,000 transconjugants from one conjugation, and transposon insertion sequencing (Tn-seq) of a pool of 1065 colonies revealed 965 unique insertion sites throughout the genome, indicating it would be feasible to generate a genome-wide mutant library using this transposon construct.

Using the pMarC9-R6k transposon, we then generated a randomly-barcoded transposon insertion mutant library in *P. sakaiensis* to enable scalable downstream screening and mutant fitness profiling (**Figure 1D**). We generated a pool of ∼4 million randomly barcoded transposons (pMarC9-RB) and delivered this library to *P. sakaiensis* by conjugation, resulting in ∼7.7 x 10^5^ transconjugants (**Supplementary Figure 2**). The genes disrupted by a transposon across the pool of mutant strains were identified and linked to their corresponding unique barcodes using a modified version of RB-TnSeq^36^ (see **Methods**). RB-TnSeq analysis identified 353,908 unique transposon barcoded strains across 44,211 unique genomic positions in the *P. sakaiensis* pooled mutant library (**Figure 1D**, **Supplementary Figure 3**). We found that 83% of all genes contained a transposon located within the central 80% of the gene, while 90% of genes larger than 600 bp had at least one insertion (**Supplementary Figure 3A-B**). We observed some insertional preference with a mean of 8.0 barcodes per insertion site, but insertions showed little positional bias across the whole genome (**Figure 1D**; **Supplementary Figure 3C**; **Supplementary Data Table 1**), indicating a library diversity sufficient for high-throughput fitness assays.

### Identifying *P. sakaiensis* genes involved in PET degradation

Using our large barcoded mutant library, we screened for *P. sakaiensis* variants with either increased or decreased fitness when growing on PET. We performed a 20-day growth experiment using either strips of amorphous PET film or a maltose control as the major carbon source (**Figure 2A**). Three replicates of the pooled *P. sakaiensis* mutant library were inoculated into YSV (yeast extract-sodium carbonate-vitamins) minimal medium supplemented with either PET film or 0.1% (w/v) maltose as the primary carbon source. Every five days over the 20-day experiment, a consistent fraction by OD of the cultures was used to seed new samples with fresh growth media to continually enrich fast-growing variants and deplete slow-growing variants through each passage. For cultures supplemented with PET film, the bacteria physically attached to the PET surface (biofilm lineage) and bacteria suspended in the liquid media (planktonic lineage) were each separately seeded into new PET cultures on day 5. Within each 5-day cultivation, we observed growth on PET ranging from maximum OD_600_ 0.21 to 0.84 in the PET planktonic lineages and OD_600_ 0.11 to 0.47 in the PET biofilm lineages (**Supplementary Figure 4**). Based on these growth measurements, we estimate that there were 11 doublings in the PET planktonic lineage, 7 doublings in the PET biofilm lineage, and 10 doublings in the maltose condition by day 20 of the experiment. The rate of degradation of the PET film in each culture was measured at each 5-day interval. PET biodegradation rates were comparable both within and between the two lineages until day 20, where a single culture from each lineage (planktonic #3 and biofilm #2) showed 2-4x increases in the biodegradation rate (**Figure 2B**).

**Figure 2.**
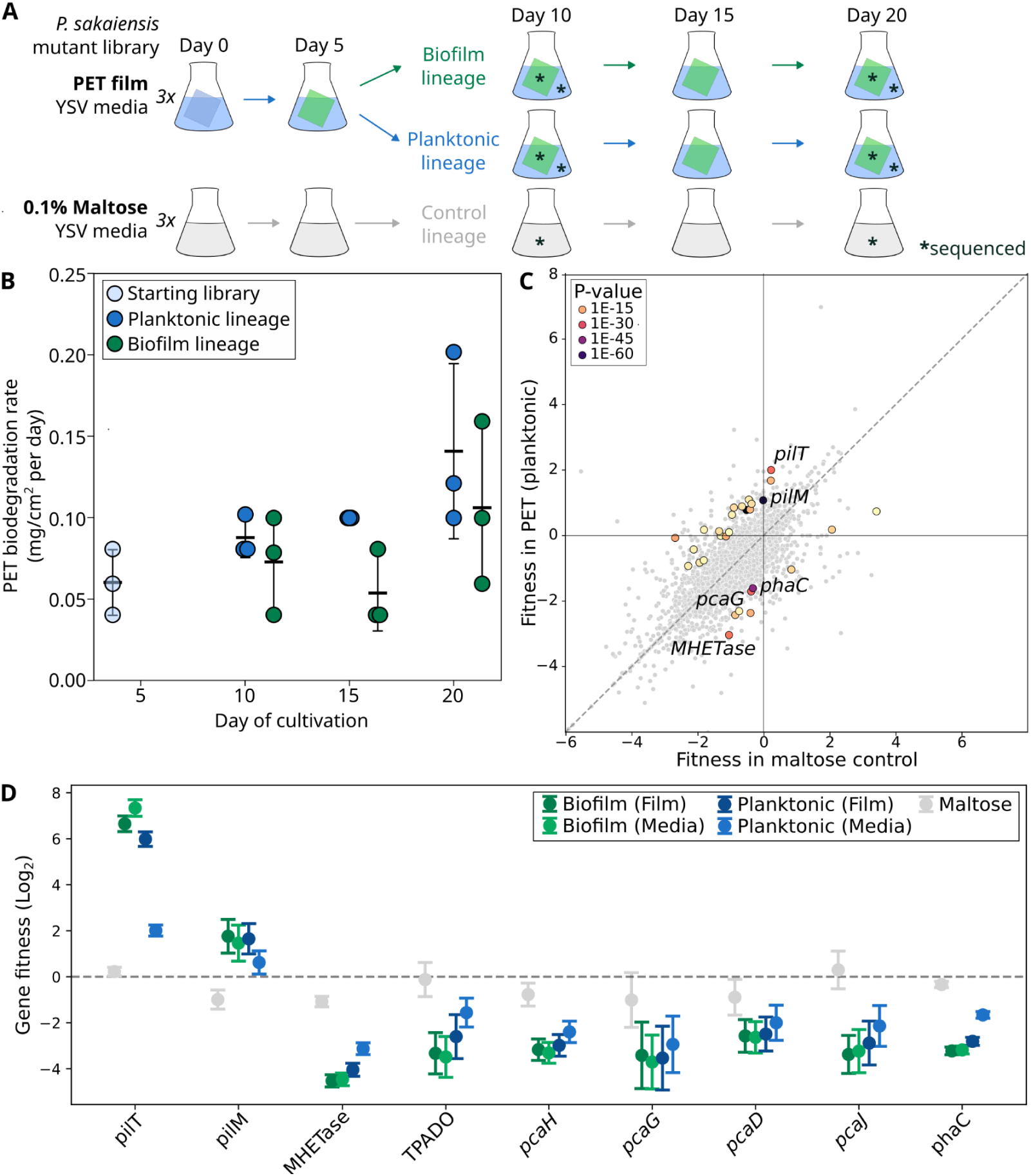
PET screen results. **(A)** Experimental timeline. *P. sakaiensis* mutant library cultures were passaged in triplicate every five days. Biofilm and planktonic fractions from both PET culture series were sequenced for RB-TnSeq analysis on day 10 and day 20. **(B)** The rate of PET degradation by PET planktonic and biofilm lineages over the 20-day experiment. Error bars represent SEM. **(C)** RB-TnSeq gene fitness in PET (planktonic) vs maltose (control) at day 10 in the experiment. Statistically significant genes are highlighted. **(D)** RB-TnSeq gene fitness of key genes predicted to be involved in PET degradation in each experimental condition. Error bars represent 99% confidence intervals.

We amplified and sequenced the barcodes of strain variants in the library under all growth conditions after 10 and 20 days of growth to quantify and compare the abundances of variants. The fitness of each gene knockout was calculated as the difference in log_2_ mean abundance of barcodes associated with that gene (i.e., strains in which that gene was knocked out) from the input library (see **Methods**). The mean number of barcoded variants detected in samples taken from PET film fractions was 68,382, lower than the mean variant number detected in the input libraries (280,867) or the maltose samples (171,365), indicating an increased degree of population bottlenecking during colonization on the PET films (**Supplementary Figure 5**). Principal component analysis of barcoded strain abundances showed an overall separation between PET and maltose cultures, indicating consistent differential selection of gene knockouts between these conditions (**Supplementary Figure 6**).

We observed 18 genes in the planktonic fraction by day 10 and 97 genes in the biofilm fractions by day 10 (68 and 128 by day 20, respectively) with statistically significant decreased fitness when growing on PET compared to the maltose controls, indicating these genes are conditionally essential for growth on PET (Mann-Whitney U; *P*<0.05; FDR=5%) (**Supplementary Data Table 2**). Of the genes conditionally essential for growth on PET, we found that mutants affecting the MHETase enzyme in the PET degradation pathway led to the greatest change in fitness (**Figure 2C**). This is consistent with biological understanding of PET degradation pathways in *P. sakaiensis,* in which secreted PETase generates MHET through hydrolysis of PET, and MHETase breaks down MHET into terephthalic acid and ethylene glycol^12,37^. While we identified seven strains with insertions in the PETase gene, and these PETase mutants had decreased fitness in PET in each condition, they were not statistically significant (Mann-Whitney U; *P*=0.16; FDR=5%). This comparison may be underpowered, given that there were few strains with PETase mutations. The insertion positions of the transposon in these strains were also located near the beginning of the PETase gene sequence, and may therefore fail to disrupt function, despite falling in the central 80% of the gene. Alternatively, as the PETase enzyme is secreted extracellularly, PETase-deficient cells may not suffer a significant disadvantage when they represent only a fraction of the population and can benefit from the overall extracellular concentration of PETase. The same dynamic would not be expected to occur for MHETase if predictions of it as a periplasmic enzyme are correct^12^. Intriguingly, a lipase chaperone gene genomically adjacent to the PETase gene, only recently recognized for its ability to improve PETase function^38^, was found to be conditionally essential for growth on PET in the biofilm samples at day 10 in the experiment (Mann-Whitney U; *P*<0.02; FDR=5%).

These data provide the first functional genetic evidence for the hypothesis that *P. sakaiensis* degrades PET through a TPA intermediate, which is then converted to protocatechuate (PCA) and subsequently degraded by a *pca* pathway^12,39^. The screen identified *pcaH*, *pcaJ*, and *pcaD* homologs that were conditionally essential for growth on PET (**Figure 2D**). The *pcaJ* and *pcaD* homologs had not been previously reported, and are not genomically colocalized with *pcaGH*. The terephthalate 1,2-dioxygenase (*TPADO*) gene downstream of the MHETase was also found to be conditionally essential for growth on PET, providing functional evidence this gene is involved in degradation of the TPA breakdown product of PET.

It has been previously noted that *P. sakaiensis* possesses an operon for biosynthesis of poly(3-hydroxybutyrate) (PHB), and that biosynthesis of PHB is increased during fermentation on PET^40^. We found that the central gene in this operon, *phaC*, is strongly conditionally essential for growth on PET (**Figure 2D**). This suggests that not only does *P. sakaiensis* produce PHB from PET, but that disruption of this biosynthetic pathway disrupts carbon metabolism and correspondingly impairs growth on PET. Therefore, production of PHB appears directly metabolically linked to growth on PET in *P. sakaiensis*.

We observed significant fitness benefits on PET for 25 gene mutants in the planktonic fraction by day 10 and 52 genes in the biofilm fraction by day 10 (43 and 144 by day 20, respectively) (Mann-Whitney U; *P*<0.05; FDR=5%) (**Supplementary Data Table 2**). These represent candidate genes that, when knocked out, may improve *P. sakaiensis*’ growth rate and/or biodegradation rate on PET substrates. The most significant (Mann-Whitney U; *P*<5.7*10^-47^; FDR=5%) of these genes was *pilT*, which encodes an ATPase enzyme that powers the retraction of type IV pili involved in motility, DNA uptake, and surface sensing (**Figure 2C-D**). In *Pseudomonas aeruginosa*, *pilT* mutants are reported to form larger biofilm structures, possibly as a consequence of their nontwitching, hyperpiliated phenotype^41,42^. A similar Δ*pilT* phenotype in *P. sakaiensis* may result in denser biofilm formation on PET leading to improved degradation of the substrate.

Fitness of mutants in the PET biofilm seeded lineages had a weak but significant correlation with the fitness of mutants in the PET planktonic seeded lineages at day 10 (R^2^=0.28; *p*<10^-6^), and more so at day 20 (R^2^=0.47; *p*<10^-13^) (**Supplementary Figure 7**). This suggests overall similar selection dynamics on the PET film and in the media; however, there were a few gene mutants with significantly higher fitness advantages on the PET film compared to the planktonic fraction. In particular, the *pilT* mutant fitness advantage was significantly higher on PET film compared to the planktonic fraction, consistent with this mutant potentially conferring advantages in dense biofilm formation (**Figure 2D**).

We tested for the impact of operonic polar effects on fitness by comparing the fitness for each gene with its adjacent upstream and unidirectional gene versus a randomly sampled gene. There was a significant correlation between fitness of genes and their upstream genes, but the variation explained by this correlation was very low (linear least squares regression; *R*^2^=0.03; *P*< 10^-22^) (**Supplementary Figure 8**). Therefore, polar effects are unlikely to strongly influence the observed fitness of the majority of genes in this dataset, similar to RB-TnSeq libraries described for *Pseudomonas stutzeri* and *Shewanella oneidensis*^36^.

### Screening the *P. sakaiensis* transposon mutant library for genes associated with genetic tractability

To identify more genetically tractable *P. sakaiensis* variants, we designed a screen to identify *P. sakaiensis* mutants with improved ability to receive DNA via conjugation (**Figure 3A**). In this screen, we delivered a replicative plasmid identified by the POSSUM screen with a pSa ORI and a gentamicin resistance marker (plasmid pGL2_170) to the transposon mutant library in a large conjugation. After 48 hours of liquid selection, we harvested DNA for sequencing and performed RB-TnSeq analysis to identify transposon mutants that were enriched. Mutants in genes which improve plasmid uptake (“conjugation activators”) would increase in abundance post-conjugation, whereas mutants of genes which decrease plasmid uptake (“conjugation inhibitors”) would decrease in abundance^43^.

**Figure 3.**
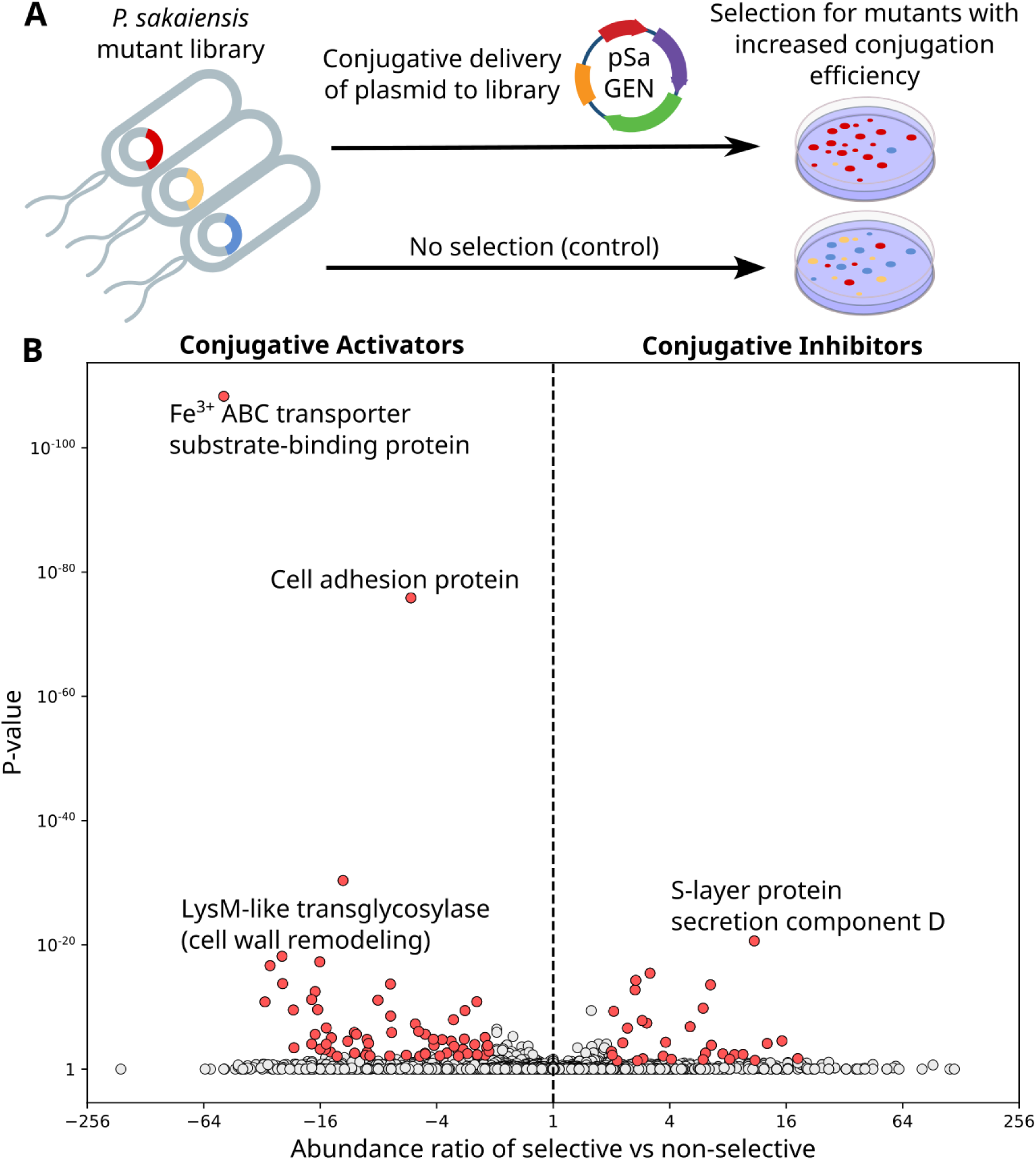
Conjugation tractability mutant screen results. **(A)** Experimental schematic. A replicative plasmid was conjugated to the transposon mutant library and transconjugants were selected for with gentamicin. Library mutants more amenable to conjugation will appear in greater frequency post-delivery when compared to a no-selection control. **(B)** Barcode abundance enrichment of gene mutants in the plasmid delivery screen. Each point represents one gene. Positive values indicate greater abundance in the selective condition, while negative values indicate lower abundance.

We identified 23 gene mutants which increased significantly after conjugation and selection that may therefore act as conjugative inhibitors (Mann-Whitney U; *P*<0.05; FDR=5%) (**Figure 3B**; **Supplementary Data Table 3**). The most significant (Mann-Whitney U; *P*<2.4*10^-18^; FDR=5%) of these was a homolog of *spsD*, a secretion component for S-layer formation. Four other genes in the same S-layer secretion and formation operon were also identified as potential conjugation inhibitors, including a remote protein homolog of *slr4*^44^, likely to form the subunit of the crystalline surface protein array (**Supplementary Data Table 3**). Bacterial S-layers are two-dimensional protein arrays on the cell surface^45^, and have been shown to be barriers to electroporation in *Paenibacillus alvei*^46^. Knocking out these genes for S-layer biogenesis in *P. sakaiensis* may therefore similarly improve conjugation efficiency and potentially electroporation as well. One of the Type I R-M systems was also identified as a conjugative inhibitor, indicating functional evidence that R-M has the potential to reduce conjugation efficiency in *P. sakaiensis* (**Supplementary Data Table 3**).

This screen further identified 56 potential conjugative activators, or gene mutants which decreased significantly after plasmid delivery by conjugation and selection (Mann-Whitney U; *P*<0.05; FDR=5%) (**Figure 3B**). These included genes involved in iron competition, cellular adhesion, and cell wall remodeling, which may facilitate interactions during conjugation. Peptidoglycan transglycosylases are known to play a role in creating local gaps and breakages in the cell wall during conjugation^47^. Finally, we note that the use of gentamicin in the selective condition and not the control implies that gene mutants that reduce gentamicin resistance^48^ may also appear as “conjugative activators” in this experiment, when in fact they are related to the antibiotic selection.

## Conclusion

This work introduces genetic engineering tools for the plastic-degrading bacteria *P. sakaiensis,* including a conjugation-based delivery toolkit comprising a donor, conjugation protocol, and an origin of transfer (*oriT*) DNA sequence as well as synthetic biology parts: a plasmid ORI, multiple selectable markers, promoters, and a transposon system. With these capabilities, we have developed a pooled genome-wide transposon mutant library and demonstrated its use to provide functional evidence for both previously hypothesized and novel metabolic functions of genes. We also demonstrated the application of this pooled mutant library to characterize factors involved in conjugative tractability of this strain. Genetic tools for *P. sakaiensis* can enable downstream experiments and engineering efforts, including *in vivo* expression of optimized PETases, functional characterization of genes involved in PET degradation, and heterologous expression of pathways for high-value products.

Our RB-TnSeq screen for fitness on PET provided the first functional evidence for the involvement of several genes in PET degradation in *P. sakaiensis*, including homologs of *pcaD*, *pcaJ*, *pcaGH*, and *TPADO*. The results of the screen also suggest that knockouts of two genes, homologs of *pilT* and *pilM*, may increase growth on PET, which could represent valuable targets in engineering *P. sakaiensis* for improved PET degradation. An important role of biofilm formation in microbial PET degradation has been previously recognized for *P. sakaiensis*^49^, and these gene mutants may increase biofilm cell density. Previously, it was reported that at least ∼9% of the carbon from PET is converted into PHB by *P. sakaiensis*^40^, and here we provide functional evidence of a metabolic dependence between PET degradation and PHB production. Further, our RB-TnSeq screen for plasmid uptake identified the *P. sakaiensis* S-layer as a potential barrier to transformation by conjugation, and S-layer knockouts could therefore be prioritized in engineering of a more tractable *P. sakaiensis* chassis, although they may also impact PET degradation phenotypes as well. Barcoded genome-wide libraries of *P. sakaiensis* will continue to be a valuable asset to the research community, as they enable rapid screening of genetic factors in other diverse phenotypes of interest as well.

There are some limitations of the high-throughput functional approaches we describe here. Polar effects have the potential to impact RB-TnSeq results^50^, although there is minimal evidence for their overall impact in our experiments. Our genome-wide library is unsaturated, with 17% of genes in the genome left unsampled, and some genes may be represented by too few insertions for sufficient statistical power when determining their fitness. The phenotypic impact of genes may also differ in a high-throughput bulk pool versus in isolation due to bet hedging and cheater effects^51^. Finally, we have not attempted isogenic knockouts of these genes, but doing so would represent a promising avenue of future research for *P. sakaiensis*.

The plastic sector is a high CO_2_ emitting industry, and its impact on the global carbon budget is projected to intensify, reaching 56 Gt CO_2_e (∼12% of the remaining global carbon budget for international climate targets)^52^. Even if renewable energy sources are used, the chemical processes involved in plastic production account for a large fraction of plastic lifecycle emissions^52^. Therefore, for plastic to become a sustainable material, it will be crucial to target and de-carbonize plastic production as well as improve plastic waste management. *P. sakaiensis* is unique among known microbes in its ability to not only degrade PET plastic, but to simultaneously convert PET into PHB, a biodegradable bioplastic. While microbial biodegradation of plastics cannot substitute for the rapid decarbonization of plastic production, engineered microbial upcycling of plastic waste presents itself as one possible and promising future component of a circular bioeconomy.

## Materials and Methods

### Strains and growth conditions

*Piscinibacter sakaiensis* 201-F6 (formerly *Ideonella sakaiensis*) received from the Leibniz Institute DSMZ, Germany (DSMZ 112585) was grown in NBRC 802 medium (casein peptone 5 g/L, peptone 5 g/L, yeast extract 2 g/L, magnesium sulfate heptahydrate 1 g/L) at 30°C, unless otherwise specified. *Escherichia coli* BW29427 was grown in LB media supplemented with diaminopimelic acid (DAP) (60 µg/mL) at 37°C. Antibiotic supplementation for all strains is noted where relevant.

### Media screen

Growth was measured using OD_600_ in a 96-well microplate with a sample volume of 150 μL, in triplicate. Measurements were taken every 20 minutes for 91 hours using the Biotek LogPhase600 plate reader (Agilent), with continuous orbital shaking at 800 RPM. To obtain growth curves, the first hour of growth was truncated, Lowess smoothing was applied, and blanking was performed by subtracting the averaged OD_600_ of media-only blank wells from the corresponding inoculated wells. A medium is considered to support growth of the strain if an OD_600_ of 0.2 is achieved in the assay.

### Antibiotic screen

Growth was measured using OD_600_ in a 96-well microplate with a sample volume of 150 μL, in duplicate. Measurements were taken every 24 hours for 48 hours using the Biotek Synergy H1 plate reader (Agilent) with orbital shaking at 800 RPM. Blanking was applied to final measurements by subtracting the averaged OD_600_ of media-only wells from the corresponding inoculated wells. An antibiotic is considered effective at a specific concentration if the strain’s OD_600_ measurement under those conditions is less than 10% of the untreated control.

### *P. sakaiensis* gDNA extraction

Genomic DNA was extracted either in low-throughput using the DNeasy Blood & Tissue Kit (Qiagen 69506) or in high-throughput using the Applied Biosystems MagMAX™ Viral/Pathogen Ultra Nucleic Acid Isolation KingFisher Kit (Thermo Fisher Scientific A42356), according to the manufacturer’s instructions. Concentrations were measured with the Qubit™ 1X dsDNA High Sensitivity Kit (Thermo Fisher Scientific Q33231).

### Whole genome sequencing and MicrobeMod analysis

*P. sakaiensis* was sequenced using both Illumina and Oxford Nanopore (R10.4.1) technologies, and basecalling was performed with Dorado v0.4. Long-read genomes were assembled using Flye^53^ with the parameter: -g 5m -nano-hq -scaffold. Assembled genomes were corrected with 2×150 bp Illumina reads using PolyPolish with default settings, and methylation patterns were identified using MicrobeMod run with default settings (Crits-Christoph et al. 2023).

### Conjugation to *P. sakaiensis*

Conjugation to *P. sakaiensis* from *E. coli* BW29427 was performed as previously described^54^ with the following modifications. Donor and recipient cultures were adjusted to OD_600_ of 100 and mixed at a 1:1 ratio (by volume, 50 µL of each). NBRC 802 1.5% agar media supplemented with DAP 60 µg/mL was used as the conjugation media. Conjugation plates were incubated for 20 hours unless otherwise stated. Transconjugants were selected on NBRC 802 1.5% agar media supplemented with the appropriate antibiotic (CAM 34 µg/mL, GEN 20 µg/mL, KAN 75 µg/mL, or TET 10 µg/mL) unless otherwise stated. Colonies were counted manually after 4 - 6 days.

### ORI screen

The ORI screen was performed as previously described using the POSSUM toolkit^31,54^ with the following modifications. Three libraries (pGL2_149, pGL2_150, pGL2_159; **Supplementary Table 4**, **Supplementary Figure 1A**) were conjugated on full petri dishes using the protocol as noted in the “Conjugation to *P. sakaiensis*” section above. Petri dish conjugations were selected using GEN 20 µg/mL, KAN 75 µg/mL, and TET 10 µg/mL on solid media. Six libraries (pGL2_217-222; **Supplementary Table 4**) were conjugated in 96-well plate format. These conjugations were selected using GEN 1.25 µg/mL, KAN 200 µg/mL, and TET 2.5 µg/mL in liquid media. The conjugative donor strain in all cases was *E. coli* BW29427.

### Tn-seq analysis of pMarC9-R6k pilot

gDNA extraction of scraped pMarC9-R6k transconjugants was performed as noted in the “*P. sakaiensis* gDNA extraction” section above.

#### Preparation of Tn-seq libraries

A 50 µL MmeI (NEB R0637; 2 U/µL) restriction enzyme digest reaction in 1x rCutSmart Buffer (NEB B6004; 10x) was prepared with 4 U of enzyme and 2 µg of input gDNA. The reaction was incubated at 37°C for 15 minutes before heat inactivation at 65°C for 20 minutes. The restriction digest was then added to an Antarctic Phosphatase (AnP, NEB M0289) reaction containing 10 µL AnP Reaction Buffer (10x), 5 µL AnP (5 U/µL), and 35 µL molecular biology grade water (MGW) for incubation at 37°C for 30 minutes and heat inactivation at 70°C. Then, the entire 100 µL reaction volume was cleaned with the DNA Clean & Concentrator-5 kit (ZYMO D4013) with the elution done in 22 µL MGW. 10 µL of the cleaned DNA was transferred to a fresh tube for 15 µL IDT pMarC9 P5 adapter duplex (1 µM; **Supplementary Table 5**) and 25 µL NEB Instant Sticky Ligase Mix (NEB M0370) to be added in the next ligation step. The reaction was incubated at room temperature for 5 minutes. DNA was again purified using the same DNA Clean & Concentrator-5 kit cleanup as before. 5 µL of DNA was then transferred to a fresh tube for a three-primer PCR amplification reaction using KAPA HiFi HotStart ReadyMix (Roche 09420398001). A 20 µL reaction was prepared using 10 µL KAPA HiFi HotStart ReadyMix (2x), 2 µL marC9_P7 primer (1 µM), 1 µL i5xx primer (10 µM), 1 µL i7xx primer (10 µM), and 1 µL MGW. The reaction went through a standard two-step thermal cycling protocol according to manufacturer’s instructions: 72°C annealing and extension temperature for 1 minute for 23 cycles. 10 µL of the PCR reaction was loaded into an E-Gel EX 2% Agarose gel (Thermo Fisher Scientific G402022) for gel extraction with the Zymoclean Gel DNA Recovery Kit (ZYMO D4001) performed according to manufacturer’s instructions.

#### Sequencing of Tn-seq libraries

The purified library was quantified using the Qubit™ 1x dsDNA High Sensitivity Kit. The rest of the library preparation was completed according to the Denature and Dilute Libraries Guide for MiniSeq System by Illumina, then loaded and sequenced on an Illumina MiniSeq in a 2 x 150 bp paired-end run.

### Construction of pMarC9-RB, a barcoded mariner transposase vector

Using pMarc9-R6k as a template, barcodes were added using N-mers from IDT primers (**Supplementary Table 5**). This PCR was done in three separate 50 µL reactions, each with 0.5 ng input template and using JL0001 and JL0002 primers, following standard KAPA HiFi HotStart (Roche 07958897001) conditions with a 64°C annealing temperature and 3 minute extension time for 24 cycles. The PCR products were pooled and cleaned with AMPure XP Beads (Beckman Coulter A63881) using a 0.5x bead to sample volume ratio and 25 µL elution volume. 3 µg of the cleaned PCR product was circularized with KLD enzyme mix (NEB M0554S) in a 30 µL reaction. The circularized product was then cleaned with the DNA Clean & Concentrator-5 kit. We first transformed the cleaned assembly into commercial electro-competent *E. coli* pir-116 cells (Biosearch Technologies EC6P095H), which have a higher transformation efficiency than our conjugative donor *E. coli* BW29427. We performed five individual 50 µL electroporations with 2.5 µL input DNA each, pooling all the transformations together after a 2 hour recovery. We performed liquid selection in 50 mL LB supplemented with CARB 100 µg/mL for ∼24 hours. We estimate a yield of ∼4 x 10^6^ barcodes from this transformation based on colony counts from a dilution series. We then prepared DNA using the QIAprep Spin Miniprep kit (Qiagen 27106) and pooled six preparations (∼90 ng/µL) to electroporate into *E. coli* BW29427. Six individual 50 µL electroporations with 2.5 µL input DNA were performed in total, with all transformations pooled together after a 2 hour recovery. We performed liquid selection in 150 mL LB supplemented with CARB 100 µg/mL for ∼24 hours. We estimate ∼9.5 x 10^7^ transformants in BW29427, exceeding the estimated number of barcodes. A chao1 estimate of the barcode diversity in the BW29427 transformant library from ∼7 million reads ranged from 3-4 x 10^6^ barcodes, corroborating the original estimate from the colony counts.

### Generation of a *P. sakaiensis* genome-wide mutant library

Conjugation was performed from a library of *E. coli* BW29427 pMarC9-RB variants to *P. sakaiensis* as described in the “Conjugation to *P. sakaiensis*” section above with the following modifications. Six individual conjugations were performed simultaneously, scraped with PBS and pooled with a combined total volume of ∼7 mL. Then, 250 µL was plated onto 24 large (245 mm) square petri dishes containing NBRC 802 1.5% agar media supplemented with KAN 75 µg/mL. Plates were incubated for 3 days until colonies appeared. The mutant library was harvested by scraping and pooling transconjugants from selective plates in a final volume of ∼20 mL. Colonies were counted using a dilution series to determine library size. The mutant library was stored in glycerol stocks for future experiments.

### RB-TnSeq library preparation and sequencing

gDNA extraction of scraped pMarC9-RB transconjugants was performed as noted in the “*P. sakaiensis* gDNA extraction” section above.

#### Preparation of RB-TnSeq libraries

A 30 µL MmeI (2 U/µL) restriction enzyme digest reaction in 1x rCutSmart Buffer (10x) was prepared with 3.6 U of enzyme and 1 µg of input gDNA. Reactions were incubated at 37°C for 15 minutes. For dephosphorylation post-digestion, a master mix of 4 µL AnP Reaction Buffer (10x), 2 µL AnP (5 U/µL), and 4 µL MGW was added directly to each sample and incubated at 37°C for 30 minutes, followed by a 20 minute heat inactivation at 70°C. DNA fragments were then purified using AMPure XP Beads according to the manufacturer’s protocol using a 1.8x bead to sample volume ratio and eluted in 10.5 µL MGW. The eluted DNA was transferred to a fresh tube for 5 µL IDT pMarC9 P5 adapter duplex (1 µM; **Supplementary Table 5**) and 15 µL Instant Sticky Ligase Mix (2x) to be added in the next ligation step. After incubation at room temperature for 5 minutes, the ligations were purified using the same 1.8x AMPure XP bead cleanup as described above, except with a 13 µL volume elution. The eluted DNA was then transferred to a fresh tube for the first PCR amplification step with KAPA Hifi HotStart PCR Kit and EvaGreen Dye (Biotium 31000, 20x). A 20 µL reaction was prepared using 4 µL KAPA Hifi Fidelity Buffer (5x), 0.6 µL KAPA dNTPs (10 mM), 0.4 µL KAPA HiFi Polymerase (1 U/µL), 1 µL EvaGreen dye (20x), 1 µL JL0003 primer (10 µM), and 0.5 µL i5xx primer (10 µM). The reaction was completed in a qTOWER^3^G thermal cycler (Analytik Jena) using a standard thermal cycling protocol according to manufacturer’s instructions: 71°C annealing temperature and 15 second extension time. The progress of each sample was tracked through the amplification curves produced in the qPCR real-time and the reaction was halted upon seeing sure signs of amplification above the reaction’s baseline - typically around 12-15 cycles. Next, 1 µL of a 10-fold dilution of the first PCR product was added to the second PCR amplification as template for a standard KAPA HiFi HotStart PCR Kit with EvaGreen Dye qPCR reaction done in a 20 µL volume exactly according to the manufacturer’s protocol using 1 µL EvaGreen dye (20x), 0.6 µL P5 primer (10 µM), and 0.6 µL i7xx primer (10 µM). The reaction was completed in a qTOWER^3^G thermal cycler using a standard thermal cycling protocol according to manufacturer’s instructions: 68°C annealing temperature and 15 second extension time. The progress of each sample was tracked through the amplification curves produced in the qPCR real-time and the reaction was halted upon seeing sure signs of amplification above the reaction’s baseline - typically around 10-13 cycles. Products were visualized on an Invitrogen E-Gel EX 2% Agarose gel to confirm library size of 288 bp and purity. A double-sided AMPure XP bead cleanup was performed according to the manufacturer’s protocol using an initial 0.4x bead to sample volume ratio and 0.8x final total bead to sample volume ratio to enrich for DNA fragments between 200-300 bp. Cleaned libraries were eluted in 12.5 µL 10 mM Tris-HCl (pH 8.0) with 0.1 mM EDTA.

#### Sequencing of RB-TnSeq libraries

DNA libraries were quantified using the Qubit™ 1x dsDNA High Sensitivity Kit and then deeply sequenced on an Illumina NovaSeq 6000 with ∼275M 2 x 150 paired-end reads.

### RB-TnSeq analysis

Quality and adapter trimming of reads was performed using BBDuk from the BBTools package^55^ v39.33 with the following settings: k=19, qtrim=r, trimq=10, maq=10, entropy=0.3. Cutadapt v5.1^56^ was used to extract genomic insertion sequences from reads with the settings: --error-rate 0.25, -g tagaccggggacttatcatccaacctgtta, --minimum-length 10, --overlap 4. Genomic sequences were mapped to the assembled *P. sakaiensis* genome using bowtie2 v2.5.4^57^ with default parameters. A custom script with the Pysam library (see **Data and Resource Availability**) was used to filter mapped reads with a MAPQ score >1 (indicating a unique mapped position) and with ≥85% identity to the reference. Barcodes associated with each read were extracted using Cutadapt to remove the surrounding primer regions, retaining just the barcode: with parameters --error-rate 0.25, -g ATCGTCACAGGTGGAGCACTTCCG, --minimum-length 15, --overlap 4, followed by parameters --error-rate 0.25, -a GCGAACATACCGACCCACCCTCAT, --minimum-length 15, --maximum-length 30, --overlap 4. Barcodes were then filtered using an approach similar to that described in Wetmore et al. 2015^36^: a barcode had to map to a unique location at least 75% of the time, the second best hit must be less than one eighth as frequent as the first hit, and had to be represented by a minimum of 3 uniquely mapped reads at that position, instead of 10. Barcodes were then linked to corresponding genes by searching for barcodes that both passed cutoff and occurred in the middle 80% of a gene; in other words, ignoring barcodes that were inserted in the first and last 10% of all basepairs of that gene.

### Biodegradation fitness screen

We performed a 20-day growth experiment with three passages of the transposon mutant library, according to the schematic described in **Figure 2A**.

#### Preparation and sterilization of PET samples

Using sterile forceps in a biosafety cabinet, previously cut pieces of amorphous PET plastic (PolymerFilms, Fresno, CA; <5% crystallinity) stored in 70% ethanol and at 4°C were removed and individually placed into separate clean weigh boats and dried for one hour. The mass of each plastic piece was recorded to the nearest 0.1 mg using an electronic scale. Plastic pieces were then UV sterilized for ∼10 minutes.

#### Preparation of culture conditions

We found that *P. sakaiensis* did not grow in YSV, a medium with very low nutrient content, in Cultivarium’s laboratory, but was reproducible in the Piedra lab. To facilitate knowledge sharing and promote reproducibility, this method is available via PRISM (prism.cultivarium.org) as an AI-assisted protocol with supporting audio, video, and text. LLM-transcribed video protocols developed by one of us (F.A.P.) are shared on growth of *P. sakaiensis* in YSV with PET (see **Data and Resource Availability**). All bacterial cultures were prepared in YSV medium^12^ (0.05% yeast extract, 0.2% ammonium sulfate, and 1% trace elements (0.1% FeSO_4_·7H_2_O, 0.1% MgSO_4_·7H_2_O, 0.01% CuSO_4_·5H_2_O, 0.01% MnSO_4_·5H_2_O, and 0.01% ZnSO_4_·7H_2_O) in 10 mM phosphate buffer, pH 7.0) supplemented with 0.2% NBRC 802 medium and with one of two different primary carbon sources: 1) two strips of amorphous PET film or 2) 0.1% maltose. Each primary carbon source was tested in triplicate in a 4-mL volume, for a total of six starting cultures. A glycerol stock of the genome-wide mutant library was thawed, washed using an equal volume of YSV medium, and then used to inoculate each culture at a starting OD_600_ of 0.05. All cultures were incubated at 30°C with shaking at 200 RPM for the duration of the experiment. At the first time point after five days, the original PET cultures were split into two separate lineages: a biofilm lineage, seeded solely from cells harvested from the biofilm that formed on the surface of the PET in the culture; and a planktonic lineage, seeded solely from cells growing in the culture media. Cultures for both lineages were set up exactly like the original amorphous PET cultures and passaged to propagate their own respective biofilm and planktonic populations, but only biofilm cells were used to seed biofilm lineage cultures and vice versa. The split between these two lineages increased the total number of active cultures in the experiment after day 5 from six to nine.

#### Sample collection and culture processing

At each time point, cultures were removed from the incubator and PET pieces were trypsinized to release the PET-adherent cells (see “Trypsinization procedure” below); planktonic cells were also harvested for optical density and absorbance readings, passaging, and sampling. OD_600_ measurements were taken of culture media using a 1 mL volume in cuvettes and a UV-Vis spectrophotometer. To estimate the OD_600_ of the biofilm samples, because the amount recovered for each culture was small, absorbances (600 nm) of both media samples and the trypsinized cells were read using 200 µL in a microplate reader. A standard curve relating the background-corrected absorbance values to the previously measured OD_600_ readings was made from all the media samples, and OD_600_ values were calculated for the biofilm samples from the standard curve. PET pieces were processed to assess for biodegradation (see “PET processing for dry mass procedure” below). Volumes needed for reseeding cultures into new 4 mL cultures at an OD_600_ of 0.05 were calculated using directly measured OD_600_ values where available. After passaging, the remaining culture volumes were pelleted by centrifugation, the supernatant discarded, and the pellet was resuspended in 1 mL YSV. For each sample type, 500 µL was aliquotted for sequencing, and 500 µL was aliquotted for cryostocks. We prepared cryostocks in 25% glycerol and froze at -80°C.

#### Trypsinization procedure

Rectangular and oval PET strips were removed from each +PET culture belonging to both biofilm and planktonic lineages and immersed in 4 mL YSV with 0.05% trypsin-EDTA and incubated at room temperature for 10 minutes. During incubation, the following was performed four times: vortex for 5 seconds, bath-sonicate for 10 seconds. Trypsinized cells were pelleted by centrifugation for 10 minutes. PET strips were carefully removed and immersed in 2% SDS to start PET processing for their dry mass. The supernatant was then removed and the pelleted cells were resuspended in 1 mL YSV. The trypsinized cells were then processed according to the “Sample collection and culture processing” section above.

#### PET processing for dry mass and biodegradation assessment

In a biosafety cabinet, the rectangular and oval PET strips were removed from trypsinization tubes, carefully blotted by touching the edge of each pair to a kimwipe, and immersed in 2% SDS. The strips were incubated at 30℃ in a shaking incubator for 30 minutes. PET strips were removed from 2% SDS, carefully blotted by touching the edge of each pair to a kimwipe, and immersed in 70% ethanol. The strips were incubated at 30℃ in a shaking incubator for 30 minutes. PET strips were removed from 70% ethanol, carefully blotted by touching the edge of each pair to a kimwipe, and each pair was placed on its own properly labeled weigh boat. Strips were left to dry for one hour in a biosafety cabinet. Each PET strip (including those from a negative control culture) was weighed and its mass recorded to the nearest mg. Observed mass changes were used to calculate PET biodegradation rates (units: mg*cm^-2^*day^-1^).

### BarSeq library preparation and sequencing

gDNA extraction of samples from day 0, 10 and 20 were performed as noted in the “*P. sakaiensis* gDNA extraction” section above.

#### Preparation of BarSeq libraries

The first PCR amplification step was a standard KAPA HiFi HotStart PCR Kit with EvaGreen Dye qPCR reaction in a 20 µL volume according to the manufacturer’s protocol using JL0003 and JL0004 primers, 500 ng input template, and 1 µL EvaGreen dye (20x) spiked in (**Supplementary Table 5**). The reaction was completed in a qTOWER^3^G thermal cycler using a standard two-step thermal cycling protocol according to manufacturer’s instructions: 72°C annealing and extension temperature for 30 seconds. The progress of each sample was tracked through the amplification curves produced in the qPCR real-time and the reaction was halted upon seeing sure signs of amplification above the reaction’s baseline - typically around 16-18 cycles. Next, 1 µL of a 20-fold dilution of the first PCR product as added to the second PCR amplification as template for another standard KAPA HiFi HotStart PCR Kit with EvaGreen Dye qPCR reaction done in a 20 µL volume exactly according to the manufacturer’s protocol using i5xx and i7xx primers and 1 µL EvaGreen dye (20x). The reaction was completed in a qTOWER^3^G thermal cycler using a standard thermal cycling protocol according to manufacturer’s instructions: 70°C annealing temperature and 15 second extension time. The progress of each sample was tracked through the amplification curves produced in the qPCR real-time and the reaction was halted upon reaching the stage of exponential amplification - typically around an Intensity [I] value of 50000 RFU and generally around 7-9 cycles. A double-sided AMPure XP bead cleanup was performed according to the manufacturer’s protocol using an initial 0.4x bead to sample volume ratio and then a 1.2x bead to sample volume ratio to enrich for DNA fragments between 150-250 bp.

#### Sequencing of BarSeq libraries

DNA libraries were quantified using the Qubit™ 1x dsDNA High Sensitivity Kit and then sequenced on an Illumina NovaSeq 6000, returning a total of ∼950M 2 x 150 paired-end reads. BarSeq libraries prepared from the “Genetic tractability screen” below returned ∼275M 2 x 150 paired-end reads.

### Analysis of BarSeq data

Reads were quality and adapter trimmed using BBDuk from the BBTools package and the parameters described above in the section “RB-TnSeq analysis”. Forward and reverse reads were then merged using BBmerge from the BBTools package^58^ with default parameters. Barcode sequences were cut from reads using Cutadapt v5.1^56^ with the parameters: -trimmed-only, -a CGGAAGTGCTCCACCTGTGACGAT, -g ATGAGGGTGGGTCGGTATGTTCGC, -n 2, --error-rate 0.25, --minimum-length 15, --overlap 10. Barcodes were matched to the database of barcodes with insertions identified by RB-TnSeq and quantified using vsearch v2.29^59^ with the parameters: --search_exact, --minseqlength 19, --maxseqlength 21. Statistical analysis of fitness results was performed using custom Python scripts. Briefly, barcode abundances within each sample were normalized to the 95% trimmed mean to account for uneven library sequencing depths and library skew. 0 values within samples were replaced with ‘pseudocounts’ of the minimum non-zero value observed across the dataset after normalization. Calculation of gene fitness was performed by taking the mean abundance of each barcode across all three replicates and then taking the mean of all barcoded strains linked to a given gene. Fitness was calculated as log_2_ of the abundances of all strains linked to a gene within a given condition minus log_2_ of the abundances of the strains linked to a gene in the three replicates of the input libraries. A Mann-Whitney U test was performed to identify statistically significant differences in both fold change and fitness values when comparing each PET condition to the maltose controls, and p-values were corrected for multiple hypotheses using the Benjamini-Hochberg Procedure^60^ (FDR=5%). Due to lower barcode diversity in samples that had been passaged or sampled on PET film, samples from all of the PET film passages and fractions on each day were combined into a single ‘biofilm’ group for Mann-Whitney U statistical analysis. Analysis of the genetic tractability BarSeq experiment (see “Genetic tractability screen” below) was performed similarly, except due to the single biological replicate in the selective condition and higher variability among the control triplicates, the median abundance of each strain was calculated across the control triplicates instead of the mean.

### Genetic tractability screen

Conjugation was performed as described in “*Conjugation to P. sakaiensis*” section above with the following modifications. Ten conjugations were performed simultaneously with pGL2_170 into the *P. sakaiensis* pMarC9-RB transposon mutant library. Twelve control conjugations were performed with a donor lacking any plasmid; the control conjugations were broken up into three replicates of four conjugations for all downstream processing. A glycerol stock of the *P. sakaiensis* mutant library was thawed on ice and inoculated (1:300 dilution) into 3x flasks containing 50 mL NBRC 802 media supplemented with KAN 75 µg/mL. The flasks were incubated at 30°C shaking at 225 RPM for 46 hours until an OD_600_ of ∼2 was reached. The pGL2_170 donor and control donor were grown to an OD_600_ of ∼1.2. We used 25 µL of donor and 50 µL of recipient, both at an OD_600_ of 100. Conjugation was performed for 2.5 hours instead of 20 hours. Conjugation plates were scraped and combined. Colonies were counted using a dilution series to determine library size. Selection was performed in 640 mL NBRC 802 media supplemented with GEN 20 µg/mL with a 6.6 mL conjugation volume inoculum. For each control, 250 mL non-selective media was inoculated with 2.5 mL conjugation volume to keep the inoculation proportions the same. Both of these cultures were incubated at 30°C shaking at 225 RPM for 54 hours before making glycerol stocks and collecting pellets for gDNA extraction.

## Data and Resource Availability

The POSSUM toolkit is available via Addgene ([POSSUM_Toolkit](https://www.addgene.org/kits/cultivarium-possum/), Kit ID #1000000234), and the pSa+GEN functional plasmid is available via Addgene ([pGL2_170](https://www.addgene.org/199102/), Plasmid #199102). Plasmid sequences used in this study are attached as Supplementary Material in GenBank format. MicrobeMod is available via [Github](https://github.com/cultivarium/MicrobeMod). Analysis code and genome data used in this work are made available at: https://github.com/cultivarium/Piscinibacter_sakaiensis_RB_TnSeq/. For inquiries on sharing the *P. sakaiensis* RB-TnSeq library, contact.

The *P. sakaiensis* genome has been deposited on NCBI with BioSample accession SAMN52661550 in BioProject PRJNA1037563. Barcode insertion sites, and gene fitness scores for RB-TnSeq experiments have been made available as **Supplementary Data Tables 1-3**.

PRISM AI-assisted protocol videos for *P. sakaiensis* growth on PET are available at: https://prism.cultivarium.org/protocols/2025-08-151755272399897 https://prism.cultivarium.org/protocols/2025-09-031756917944237 https://prism.cultivarium.org/protocols/2025-09-181758221307298

## Author Contributions

C.G., N.O., H.H.L. conceived of and designed the project. V.A.S. highlighted the need for genetic tools for *P. sakaiensis*. V.A.S. demonstrated cultivation and selection, and performed initial POSSUM experiments with S.L.B.. A.E. performed the media, antibiotic, and POSSUM ORI screens in liquid media. A.C.C. and J.L. designed the transposon experiments. J.L. built and delivered the pMarC9-RB transposon library, performed DNA sequencing workflows, and performed the genetic tractability screen. F.A.P. performed the PET degradation screen. A.C.C. processed the sequencing data and performed analysis. K.M. developed pipelines for analysis of growth, antibiotic, and ORI data. M.B.A. and all other authors assisted with writing the manuscript.

## Acknowledgements

We thank all members of the Cultivarium team for discussions throughout this project. Cultivarium acknowledges support from Eric and Wendy Schmidt as a Convergent Research Focused Research Organization (FRO).

## bioRxiv usage

As with previous preprints led by Cultivarium, a Focused Research Organization, this bioRxiv is intended as a final public manuscript, and this work has not been submitted to a journal. Comments and feedback are encouraged in the sections provided by bioRxiv.

## Supplementary Figures

**Supplementary Figure 1.**
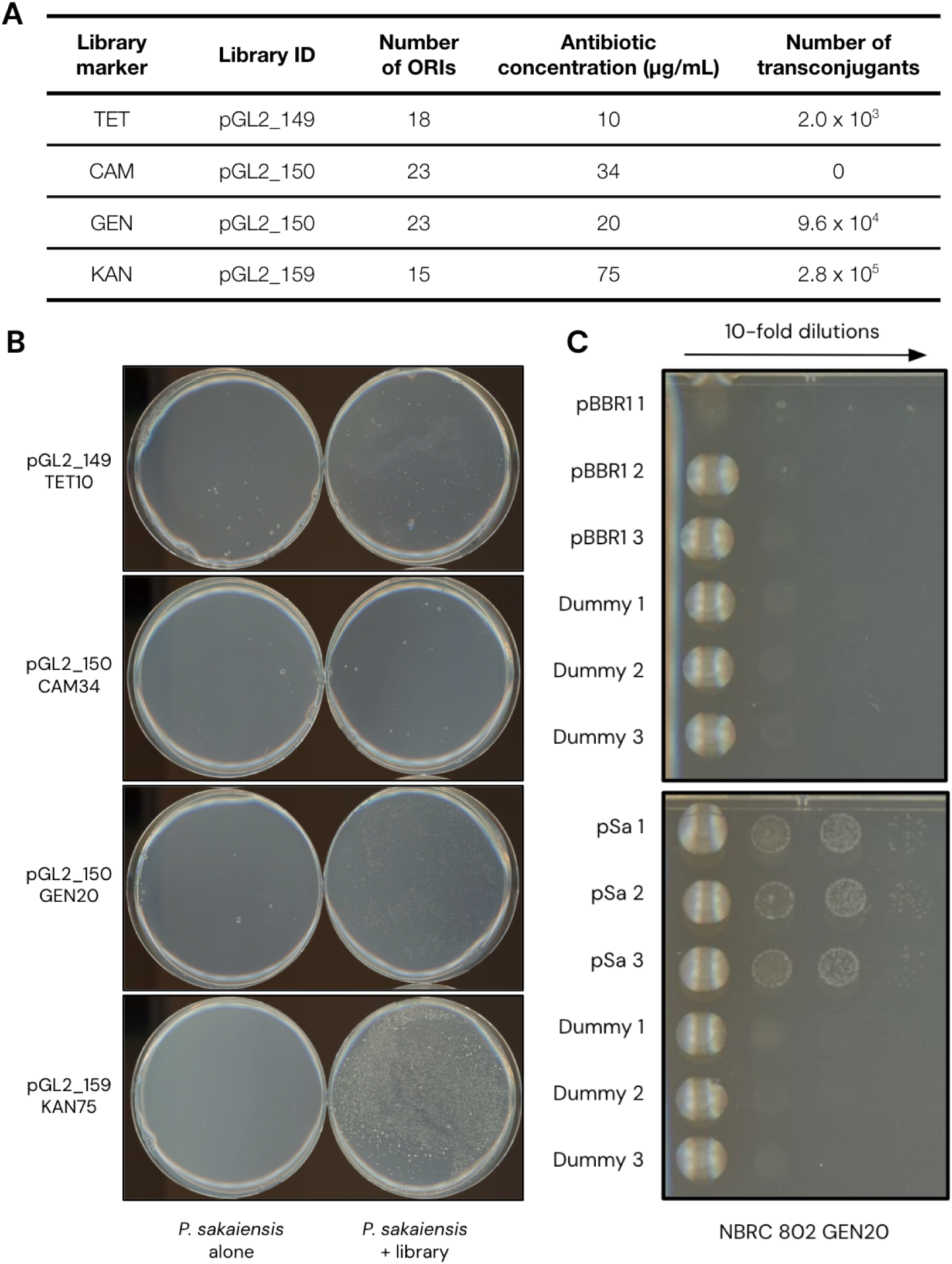
ORI-marker screen in *P. sakaiensis*. **(A)** POSSUM libraries delivered to *P. sakaiensis* in low throughput and the number of transconjugants obtained. **(B)** Selective plates following conjugation of ORI-marker libraries to *P. sakaiensis* in low throughput imaged after 6 days of growth. **(C)** Selective plates following conjugation of individual plasmids harboring pBBR1 or pSa ORI to *P. sakaiensis* compared to the non-functional dummy ORI plasmid.

**Supplementary Figure 2.**
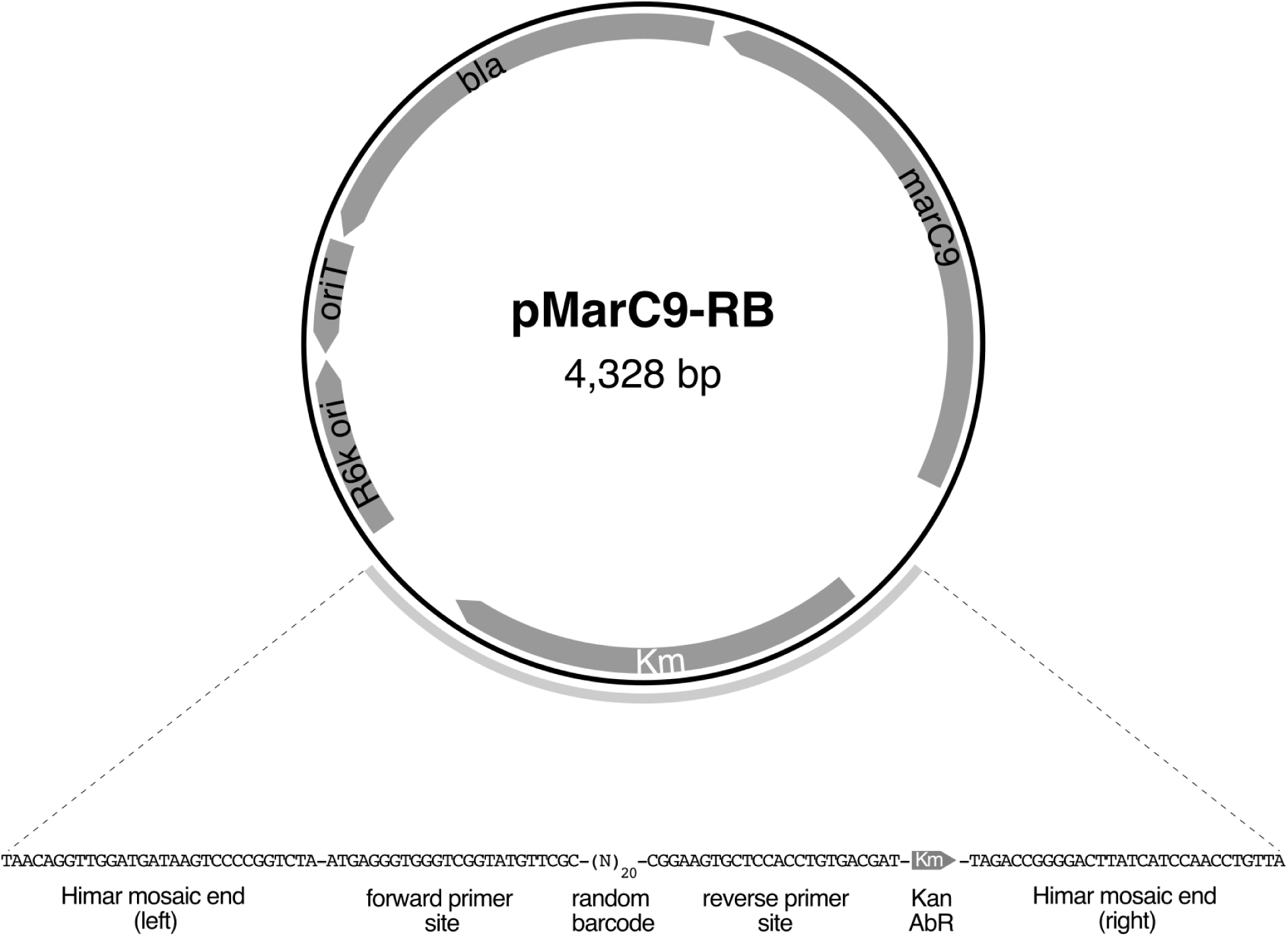
Sequence schematic of pMarC9-RB plasmid and transposon construct.

**Supplementary Figure 3.**
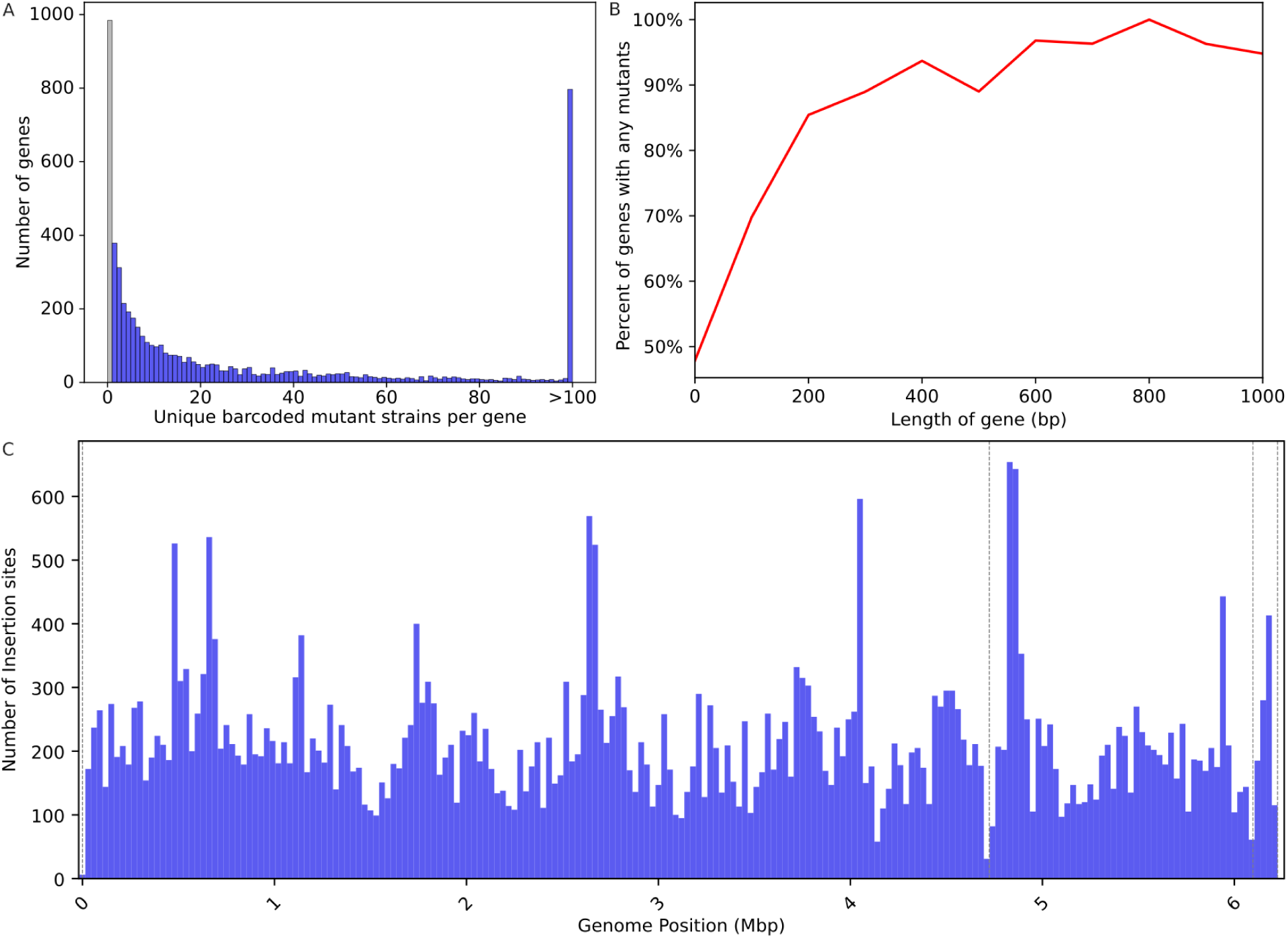
*P. sakaiensis* RB-TnSeq library quality. **(A)** Number of mutant strains with unique barcodes per gene in *P. sakaiensis*. Genes with 0 barcodes (18%) are colored in grey. **(B)** Percentage of genes with at least one barcoded mutant by length of associated gene. **(C)** Insertion site frequency along the *P. sakaiensis* genome, in 30 Kb windows. Genomic contig boundaries are denoted by grey dotted lines.

**Supplementary Figure 4.**
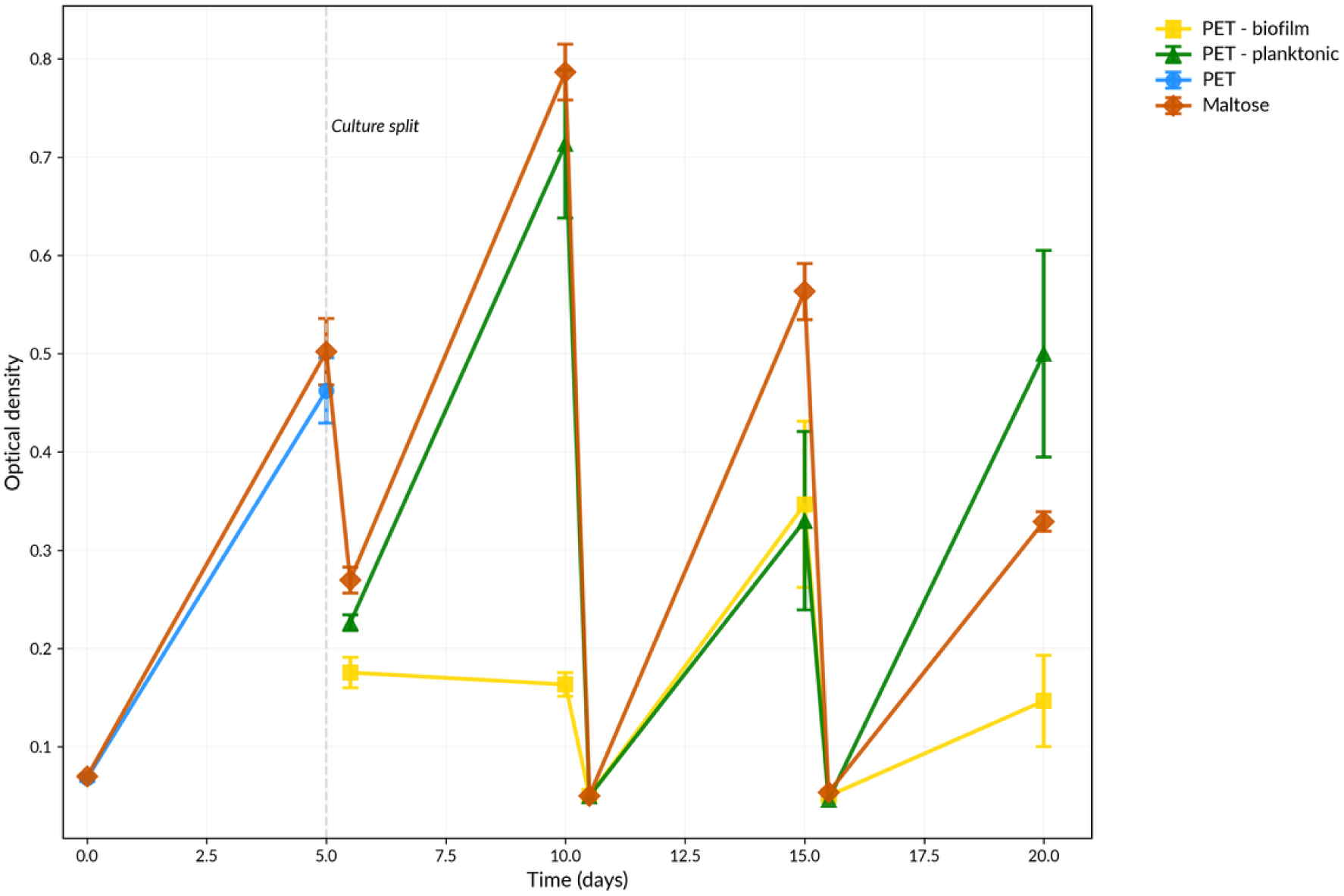
*P. sakaiensis* growth in YSV media supplemented with PET film or 0.1% (w/v) maltose as the carbon source. Growth was measured as optical density at 600 nm.

**Supplementary Figure 5.**
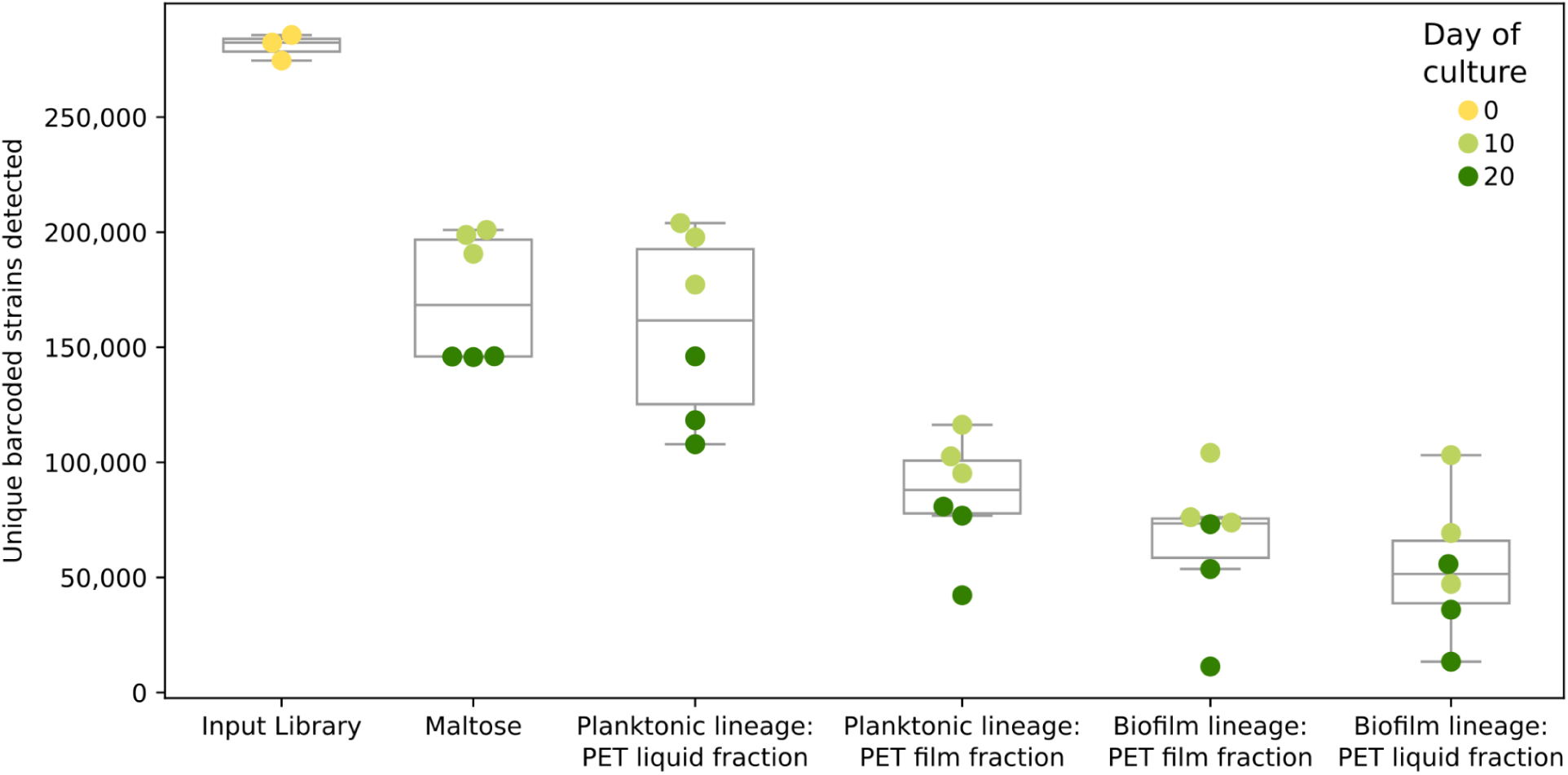
Number of detectable barcodes per sample by timepoint in the experiment.

**Supplementary Figure 6.**
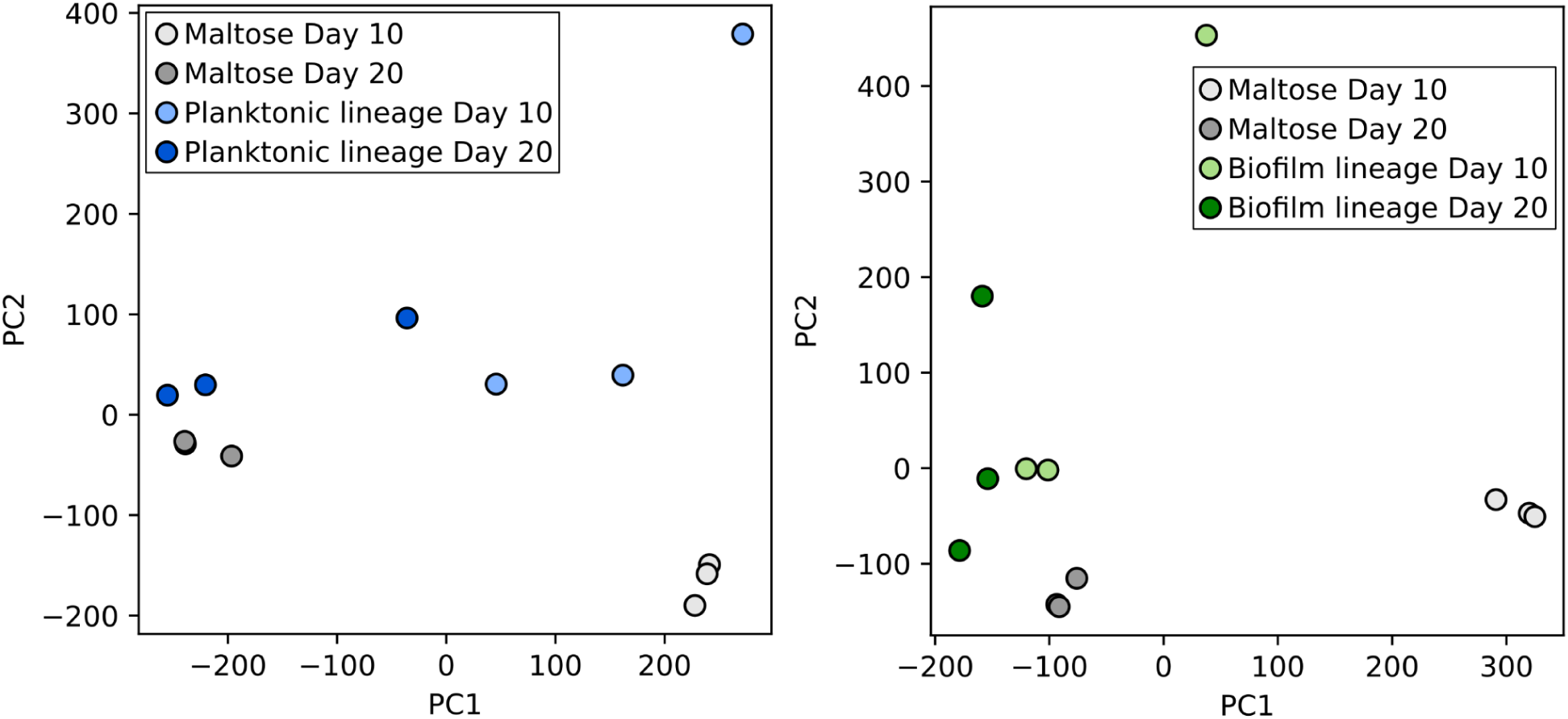
Principle component analysis of library abundances of gene mutant barcoded strains. The planktonic fraction of planktonic lineage samples (left) and the biofilm fraction of biofilm lineages (right) are compared to maltose control samples after 10 and 20 days of cultivation.

**Supplementary Figure 7.**
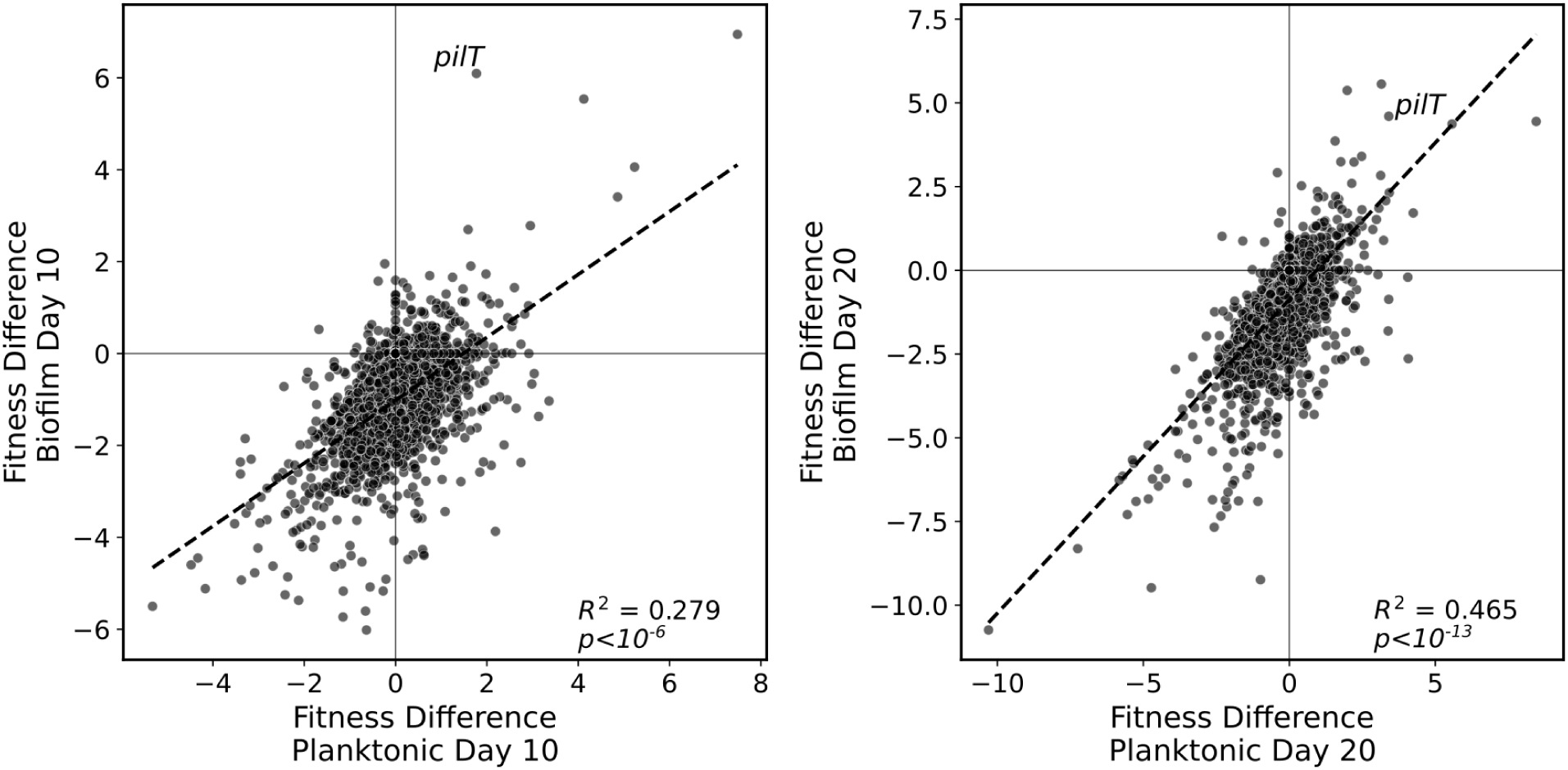
Correlation between fitness in the planktonic and film fractions.

**Supplementary Figure 8.**
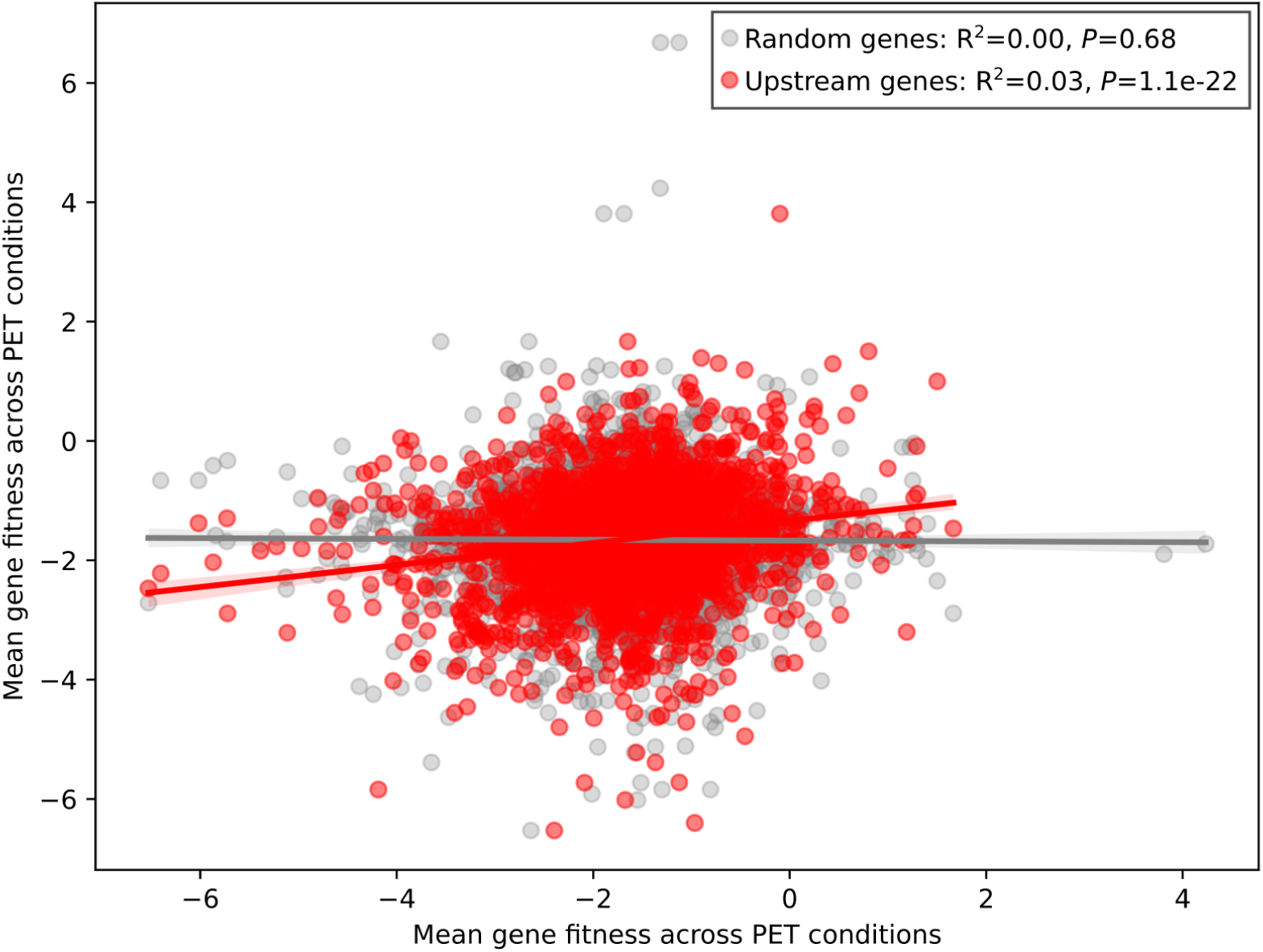
Test for polar effects between operonic genes. Each red point represents the mean gene fitness of a gene across all PET conditions compared to the fitness of its upstream gene in the same direction, if one exists. A control of gene-gene fitness comparisons across randomly sampled genes is also shown in grey. Linear least-squares regression lines for each are shown.

## Supplementary Tables

**Supplementary Table 1.** Media screen of *P. sakaiensis*. Area under the curve (AUC) is a summary variable of growth lag, yield, and rate that is calculated by integrating the growth curve over the first 24 hours.

| Status | Media | Max OD600 | Doubling time (hr) | Time to OD 0.2 (hr) | AUC |
| --- | --- | --- | --- | --- | --- |
| positive | Horikoshi-1 pH 8<sup>†</sup> | 0.998 | 7.498 | 43.889 | 0.35 |
| positive | Yeast Extract Malt Extract Dextrose pH 8<sup>†</sup> | 0.956 | 5.23 | 36 | 0.33 |
| positive | NBRC 802 | 0.84 | 6.024 | 31.778 | 0.44 |
| positive | Tryptic Soy Broth (TSB) | 0.388 | 4.76 | 71.001 | 0.13 |
| positive | Nutrient Broth pH 8<sup>†</sup> (NB) | 0.369 | - | 32.445 | 0.21 |
| negative | Halophile Basic Medium<sup>†</sup> | 0.093 | - | - | 0.045 |
| negative | R2A | 0.058 | - | - | 0.034 |
| negative | Marine Broth | 0.006 | - | - | 0.0045 |
| negative | Yeast Extract Peptone Dextrose | 0.014 | - | - | 0.032 |
| negative | Yeast Extract Nutrient Broth | 0.009 | - | - | 0.0023 |
| negative | LB | 0.008 | - | - | 0.0024 |
| negative | Halophile Complete Medium 15<sup>†</sup> | 0.011 | - | - | 0.0026 |
| negative | Brain-Heart Infusion Broth | 0.007 | - | - | 0.0018 |

<sup>†</sup>Media also contains Trace Elements (Teknova T1001) at 1X final concentration.

**Supplementary Table 2.** Antibiotic screen of *P. sakaiensis* in liquid media. MIC90 refers to the Minimum Inhibitory Concentration at which 90% of bacteria are inhibited from growing by the antibiotic. Cultures were grown in NBRC 802 media for 48 hours.

| Antibiotic | Resistant | MIC90<br>($\mu\text{g/mL}$) | Tested range<br>($\mu\text{g/mL}$) |
| --- | --- | --- | --- |
| Erythromycin | No | 4 | 0.25 - 16 |
| Tetracycline | No | 2.5 | 0.63 - 40 |
| Spectinomycin | No | $\leq 3.12$ | 3.12 - 200 |
| Chloramphenicol | No | $\leq 2.12$ | 2.12 - 136 |
| Kanamycin | No | 200 | 3.12 - 200 |
| Gentamicin | No | $\leq 1.25$ | 1.25 - 80 |
| Carbenicillin | Yes | - | 6.25 - 400 |

**Supplementary Table 3.**
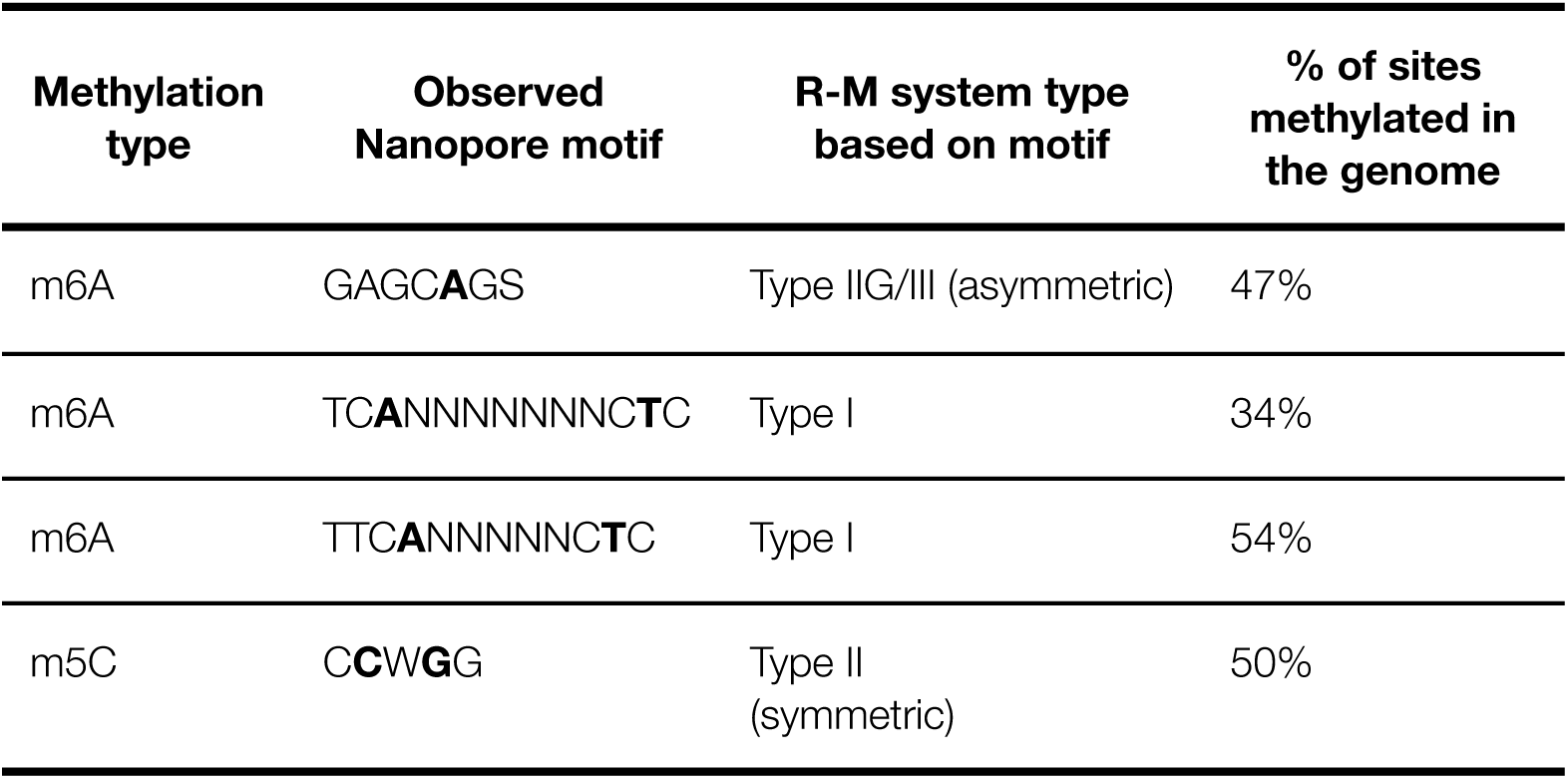
Active methylation motifs of *P. sakaiensis* identified using Nanopore sequencing and analysis with MicrobeMod. Bolded bases indicate methylated base positions in the sequence motif.

**Supplementary Table 4.**
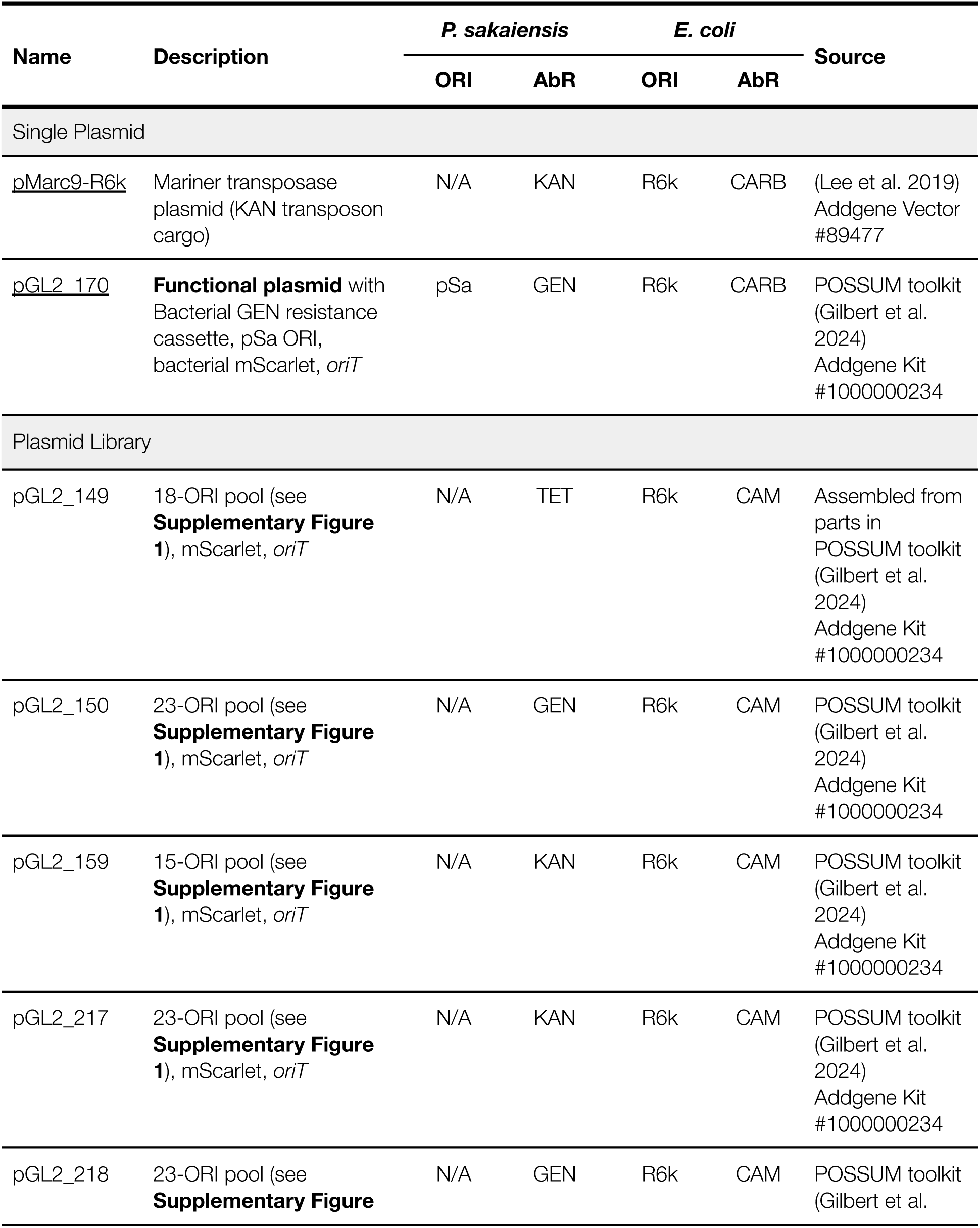
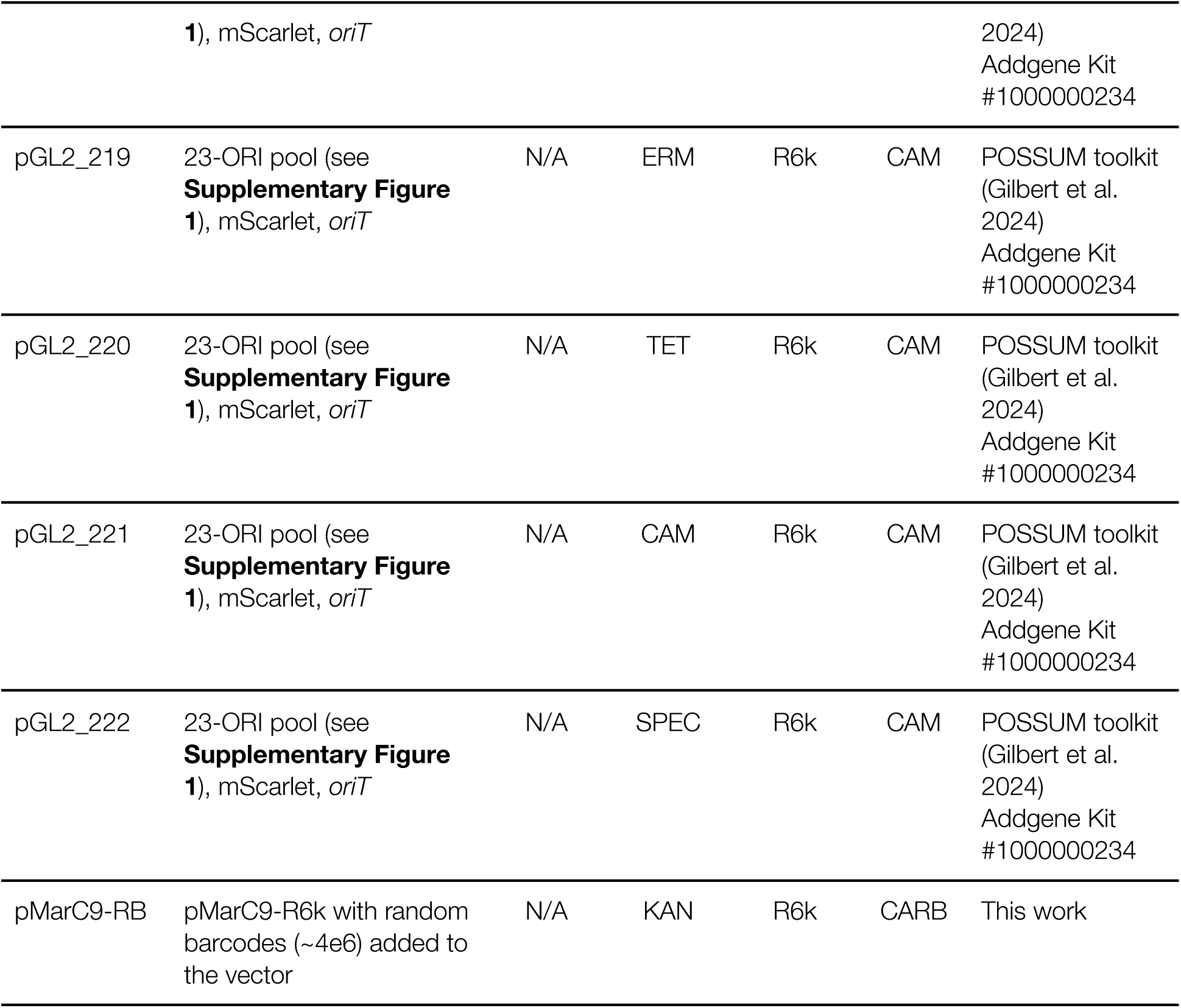
Plasmids used in this work.

**Supplementary Table 5.**
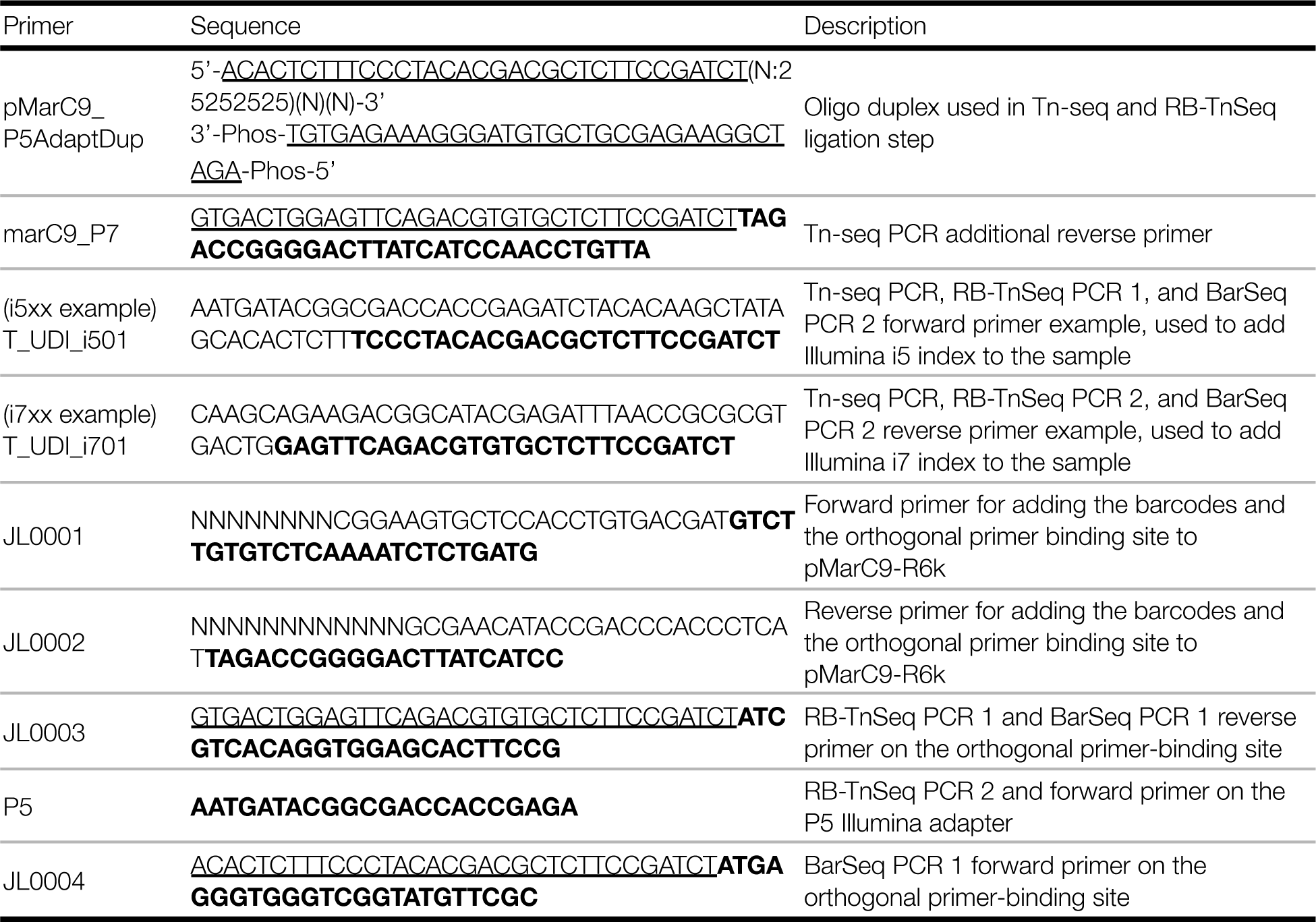
Primers used in this work. Regions of homology to the fragment being amplified are shown in bold. The underlined regions represent 5’ primer extensions which introduce stub sequences enabling binding of dual indexed Illumina i5 and i7 primers in the subsequent PCR step.

## External Supplementary Material

**Supplementary Data Table S1:** Full list of random barcode sequences and their genomic insertion sites in the *P. sakaiensis* RB-TnSeq library.

**Supplementary Data Table S2:** Abundance, fitness, and significance values for genes during the PET growth experiment.

**Supplementary Data Table S3:** Abundance, fold-change, and significance values for genes during the conjugative delivery experiment.

